# A pair of RNA binding proteins inhibit ion transporter expression to maintain lifespan

**DOI:** 10.1101/2023.05.10.540279

**Authors:** Rebekah Napier-Jameson, Olivia Marx, Adam Norris

## Abstract

Regulation of lifespan by transcription factors has been well established. More recently a role for RNA binding proteins (RBPs) in regulating lifespan has also emerged. In both cases, a major challenge is to determine which regulatory targets are functionally responsible for the observed lifespan phenotype. We recently identified a pair of RBPs, *exc-7/ELAVL* and *mbl-1/Muscleblind*, which display synthetic (non-additive) lifespan defects: single mutants do not affect lifespan, but *exc-7; mbl-1* double mutants have strongly reduced lifespan. Such a strong synthetic phenotype represented an opportunity to use transcriptomics to search for potential causative targets that are synthetically regulated. Focus on such genes would allow us to narrow our target search by ignoring the hundreds of genes altered only in single mutants, and provide a shortlist of synthetically-regulated candidate targets that might be responsible for the double mutant phenotype. We identified a small handful of genes synthetically dysregulated in double mutants and systematically tested each candidate gene for functional contribution to the *exc-7; mbl-1* lifespan phenotype. We identified one such gene, the ion transporter *nhx-6*, which is highly upregulated in double mutants. Overexpression of *nhx-6* causes reduced lifespan, and deletion of *nhx-6* in an *exc-7; mbl-1* background partially restores both lifespan and healthspan. Together, these results reveal that a pair of RBPs mediate lifespan in part by inhibiting expression of an ion transporter, and provide a template for how synthetic phenotypes (including lifespan) can be dissected at the transcriptomic level to reveal potential causative genes.

## INTRODUCTION

Aging in most organisms is associated with progressive functional impairment and susceptibility to death. The rate of aging is partially controlled by genetic pathways, including a role for transcription factors that regulate gene expression levels. For example, in the nematode worm *Caenorhabditis elegans,* it is well established that lifespan-extending treatments such as dietary restriction and reduced Insulin/IGF signaling require the activity of transcription factors including *daf-16*/*FOXO*, *skn-1/NRF,* and *hsf-1/HSF1*(1–3).

In contrast, genetic control of the aging process at the level of post-transcriptional RNA regulation has only recently come to light. In *C. elegans*, the RNA binding proteins (RBPs) *hrpu-1/HNRNPU* and *sfa-1/SF1* were both recently shown to be required for lifespan extension mediated by dietary restriction, and to regulate alternative mRNA processing (4,5). These findings, coupled with correlative data indicating that RBP expression and post-transcriptional RNA regulation are associated with lifespan in various organisms (6,7), suggest that post-transcriptional control by RBPs plays an important role in determining lifespan.

A major challenging question in the case of both transcriptional and post-transcriptional regulation of the aging process is: which of the dozens or hundreds of target transcripts are responsible for the activity of a given regulatory factor? For example, two decades of intensive efforts have gone into identifying and investigating the transcriptional targets of the DAF-16 transcription factor (8–11). Nevertheless, we still lack a complete picture of which are the causative DAF-16 targets responsible for mediating increased lifespan in response to reduced Insulin/IGF signaling (12).

We recently identified a pair of RBPs, *exc-7/ELAVL* and *mbl-1/Muscleblind*, that are combinatorially required to maintain normal *C. elegans* lifespan (13). Loss of either RBP on its own causes minimal change in lifespan, but simultaneous loss of both RBPs causes drastically shortened lifespan. We hypothesize that since the lifespan phenotype is synthetic, *i.e.* not an addition of the phenotypes of the single mutants, there might be corresponding synthetic effects on the expression of specific genes. Focus on any such synthetically-regulated target genes would allow us to narrow our target search by ignoring the hundreds of genes altered only in single mutants, and provide a shortlist of candidate targets that might be responsible for the *exc-7; mbl-1* double mutant phenotype.

Guided by this hypothesis, here we analyze the transcriptomes of *exc-7; mbl-1* double mutants and the constituent single mutants, to determine the subset of genes that are uniquely dysregulated in double mutants (more severe dysregulation than in the constituent single mutants). This filtering narrowed our focus from 687 dysregulated genes to a shortlist of 8 genes dysregulated at the level of transcript abundance. We systematically tested these candidates for functional contribution to the *exc-7; mbl-1* lifespan phenotype, and identified one such causal gene, the ion transporter *nhx-6*. *nhx-6* is highly overexpressed in double mutants compared to single mutant or wild type animals. Transgenic overexpression of *nhx-6* causes a reduction in lifespan, and deletion of *nhx-6* in the context of an *exc-7; mbl-1* double mutant partially rescues defects in both lifespan and healthspan. Together, these results reveal that the RBPs EXC-7 and MBL-1 mediate lifespan in part by inhibiting expression of an ion transporter, and provide a template for how synthetic genetic phenotypes can be dissected at the transcriptomic level to reveal potential causative genes.

## RESULTS

### *exc-7; mbl-1* double mutants have reduced lifespan and healthspan

We previously identified a genetic interaction between the conserved RBPs *exc-7/ELAVL* and *mbl-1/MBNL* resulting in severe reduction of lifespan in *C. elegans* (13). Mutations in either *exc-7* or *mbl-1* alone have little effect on lifespan, but double mutant *exc-7; mbl-1* animals have strong lifespan deficits (Figure 1A), indicating a synthetic or non-additive genetic interaction between *exc-7* and *mbl-1*. The lifespan phenotype of *exc-7; mbl-1* double mutant can be rescued by transgenic over-expression of *exc-7* or *mbl-1* (Figure 1B), showing that the severely shortened lifespan of *exc-7; mbl-1* double mutants is due to an on-target synthetic genetic interaction affecting lifespan.

The reduced lifespan of *exc-7; mbl-1* double mutants could stem from a variety of underlying processes, including accelerated aging, developmental defects, or tissue-specific defects that cause lethality. To test the possibility of accelerated aging, we examined a suite of phenotypes that undergo progressive deterioration in wild-type worms as they age. These phenotypes, including pharyngeal pumping (eating behavior), locomotion, and intestinal permeability, are often considered markers of worm healthspan, the period of an organism’s life that it is healthy (14,15).

We tested whether *exc-7; mbl-1* double mutants exhibit reductions in healthspan phenotypes, and in particular whether these phenotypes decline unusually rapidly with age. We found that the rate of pharyngeal pumping slowly declines with age in wild-type and single-mutant worms (Figure 1C and S1). *exc-7; mbl-1* double mutants have normal rates of pumping throughout development and early adulthood, but rates decline sharply thereafter (Figure 1C). We found a similar trend with intestinal barrier dysfunction, which can be revealed by the “SMURF” assay in which blue food dye seeps through faulty intestinal barriers, thus coloring the animal blue (15) (Figure 1D-E). Both wild-type and double mutant animals exhibit progressive intestinal barrier dysfunction. Double mutant animals begin adulthood with normal intestinal barrier function, but decline substantially faster than wild type (Figure 1E). Finally, a similar trend holds with locomotion (Figure 1F). Initially the locomotion of double mutant worms is indistinguishable from wild-type or single mutants, but declines sharply in adulthood (Figure 1F, S1).

In each of these cases, double-mutant worms undergoing early larval development are indistinguishable from wild type, but rapidly deteriorate with age in adult worms. These results mirror the lifespan results, in which *exc-7; mbl-1* mutant worms die rapidly upon reaching adulthood, but not during the course of development (Figure 1A-B). Based on these results we conclude that *exc-7; mbl-1* double mutants develop normally without exhibiting obvious developmental defects. Then as adults these worms experience an accelerated decline of healthspan and lifespan.

**Figure 1:**
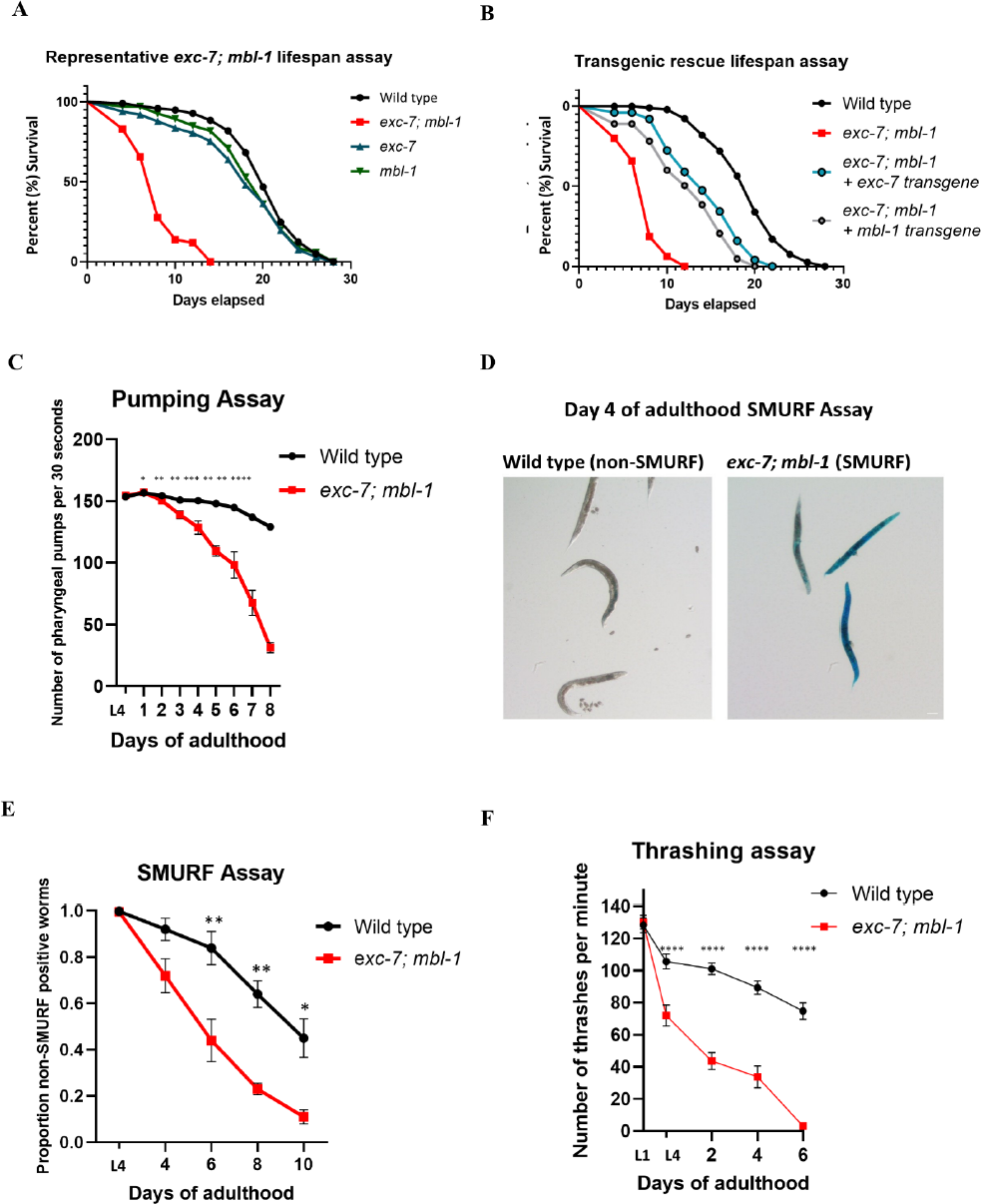
*exc-7; mbl-1* double mutants have lifespan and healthspan defects. (A) Representative *exc-7; mbl-1, exc-7*, and *mbl-1* lifespan curve. Additional biological replicates are in Figure S1. Mutations in either *exc-7* or *mbl-1* alone have little effect on lifespan, but double mutant *exc-7; mbl-1* animals have strong lifespan deficits. Log-rank test. WT (wild type). (B) Representative *exc-7* and *mbl-1* overexpression lifespan assay. The lifespan phenotype of *exc-7; mbl-1* double mutant can be rescued by transgenic over-expression of *exc-7* or *mbl-1.* Log-rank test. (C) *exc-7; mbl-1* worms start with a similar pumping rate to wild type worms as L4s but declines with age. T-test. (D) Micrographs of Day 4 adult wild type worms showing non-SMURF phenotype and Day 4 adult *exc-7; mbl-1* worms showing SMURF phenotype. Scale bar ∼100μm. (E) SMURF/ intestinal permeability assay. *exc-7; mbl-1* double mutants have increased permeability compared to wild type worms. Welch’s t-test. (F) *exc-7; mbl-1* thrashing assay. Number of thrashes per minute (wave initiation rate) from L1-D6. Initially the locomotion of double mutant worms is indistinguishable from wild type, but declines sharply in adulthood. Welch’s t-test. P-values: L1: (ns), L4: wild type vs *exc-7; mbl-*1****, D2-6: wild type vs *exc-7; mbl-1*. For all panels, ns= P>0.05, *= P≤0.05, **= P≤0.01, ***= P≤0.001, ****=P≤0.0001.

### Neuronal expression of *exc-7* and *mbl-1* controls lifespan

Expression of both *exc-7* and *mbl-1* have previously been observed in specific neuronal cell types, and *exc-7* expression has also been observed in additional tissues including muscles (16–18). We likewise observe expression of both factors in the nervous system using transgenic translational reporters under the native promoters for *exc-7* and *mbl-1* (Figure 2A). In the case of *mbl-1*, we only detect expression in the nervous system, while for *exc-7* we observe expression both the nervous system and in other tissues (Figure 2A).

To test which tissues are most important for the lifespan phenotype of the double mutant, we re-expressed either *exc-7* or *mbl-1* under tissue-specific promoters in an *exc-7; mbl-1* double mutant background. We found that for both factors, the nervous system is the primary tissue in which they are required for lifespan. Re-expression of either *exc-7* or *mbl-1* pan-neuronally leads to a strong increase in double-mutant lifespan, whereas re-expression in the intestine or in muscle results in mild or no increase in lifespan (Figure 2B-E). Together these results indicate that *exc-7* and *mbl-1* are both required in the nervous system to maintain proper lifespan.

**Figure 2:**
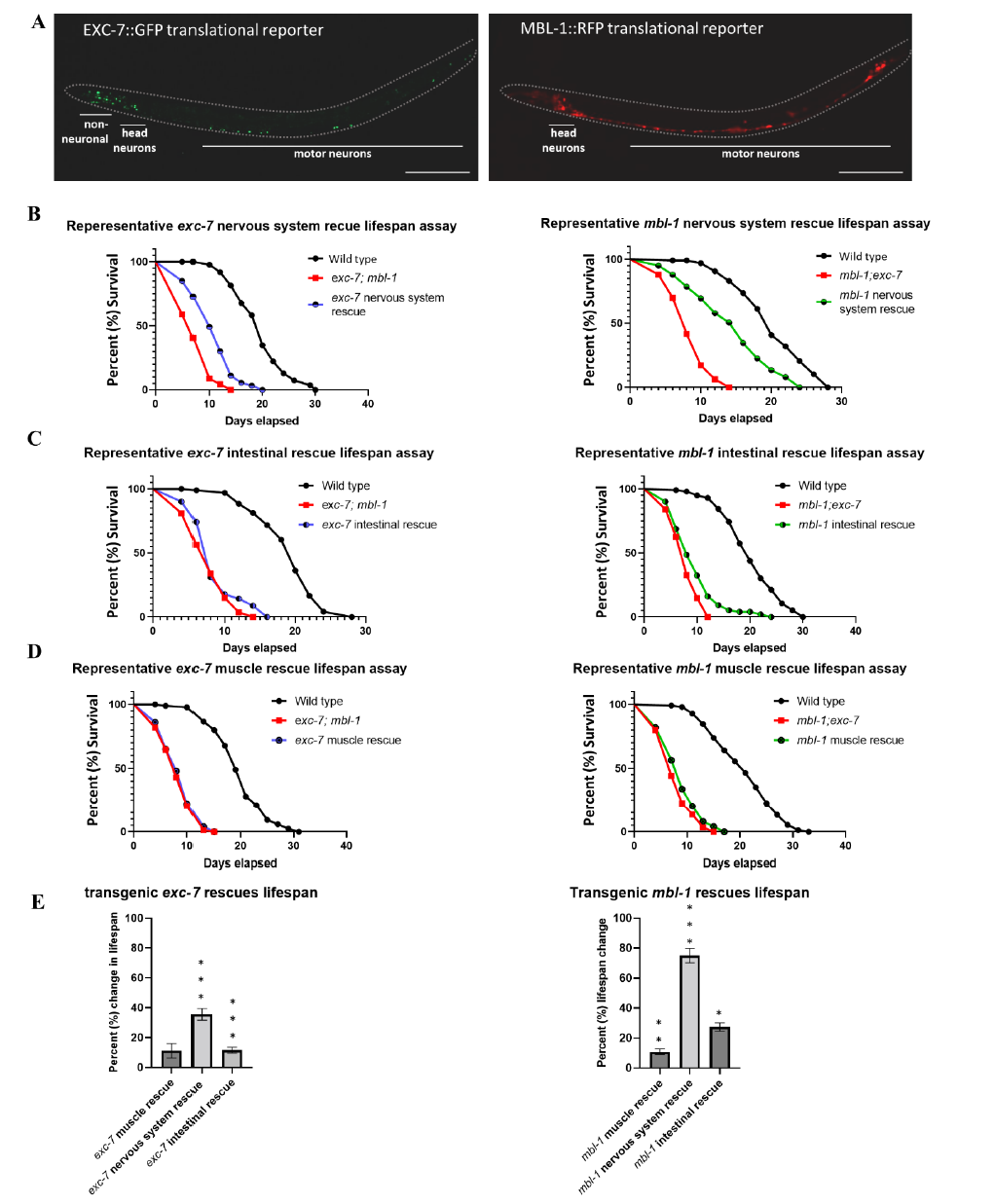
The nervous system is the primary tissue responsible for the lifespan phenotype in terms of *exc-7* and *mbl-1* expression. (A) Representative images of EXC-7::GFP and MBL-1::RFP translationally-tagged proteins driven by their native promoters. Scale bar= 100 μm. (B) Representative nervous system rescue lifespan assays. Additional biological replicates in Figure S2. Bonferroni P-values: *exc-7* nervous system rescue: exc*-7; mbl-1* vs *exc-7* nervous system rescue=****, *mbl-1* nervous system rescue: *exc-7; mbl-1* vs *mbl-1* nervous system rescue****(log-rank test. (C) Representative intestinal rescue lifespan assays. Bonferroni P-values: *exc-7* intestinal rescue: *exc-7; mbl-1* vs *exc-7* intestinal rescue=ns. *mbl-1* intestinal rescue: *exc-7; mbl-1* vs *mbl-1* intestinal rescue**. (log-rank test). (D) Representative muscle rescue lifespan assays. *exc-7* muscle rescue: Bonferroni P-values: *exc-7; mbl-1* vs *exc-7* muscle rescue(ns). *mbl-1* muscle rescue: *exc-7; mbl-1* vs *mbl-1* muscle rescue(ns) (log-rank test). (E) Summary of tissue specific rescues. Re-expression of either *exc-7* or *mbl-1* pan-neuronally leads to an increase in double-mutant lifespan, whereas re-expression in the intestine or in muscle results in mild or no increase in lifespan. P values: exc-7; mbl-1 vs *mbl-1 muscle rescue*\*\*, exc-7; *mbl-1* vs *mbl-1* nervous system rescue***, *mbl-1* intestinal rescue*, *exc-7; mbl-1* vs *exc-7* muscle rescue(ns), *exc-7; mbl-1* vs *exc-7* nervous system rescue***, *exc-7; mbl-1* vs *exc-7* intestinal rescue*** (Welch’s t test). UAS-GAL4 system was employed in order to drive expression of either *exc-7* or *mbl-1* in specific tissue. For all panels, ns= P>0.05, *= P≤0.05, **= P≤0.01, ***= P≤0.001, ****=P≤0.0001.

### Identification of transcriptomic changes unique to *exc-7; mbl-1* double mutants

Both *mbl-1* and *exc-7* have previously been shown to affect gene expression and RNA splicing (13,19,20). Therefore, to begin to probe the molecular mechanisms underlying the *exc-7; mbl-1* lifespan phenotype, we performed whole-animal RNA Seq to test for transcriptomic dysregulation in double mutants (Figure 3A). Since the lifespan phenotype of *exc-7; mbl-1* double mutants is synthetic, *i.e.* not an addition of the phenotypes of the single mutants, we hypothesized that there might be corresponding synthetic effects on the expression of specific genes. Such cases of synthetically-regulated target genes would represent good candidates for a mechanistic basis for the *exc-7; mbl-1* phenotype. We reasoned that this might circumvent a problem commonly encountered when searching for physiologically-relevant targets of gene regulatory factors: which of the dozens or hundreds of targets might be responsible for the phenotype? In our case, by focusing on transcriptomic dysregulation unique to double mutants, we can filter out the hundreds of targets dysregulated in either *exc-7* or *mbl-1* single mutants, and focus on a smaller set of targets that are uniquely dysregulated in double mutants (Figure 3B).

**Figure 3:**
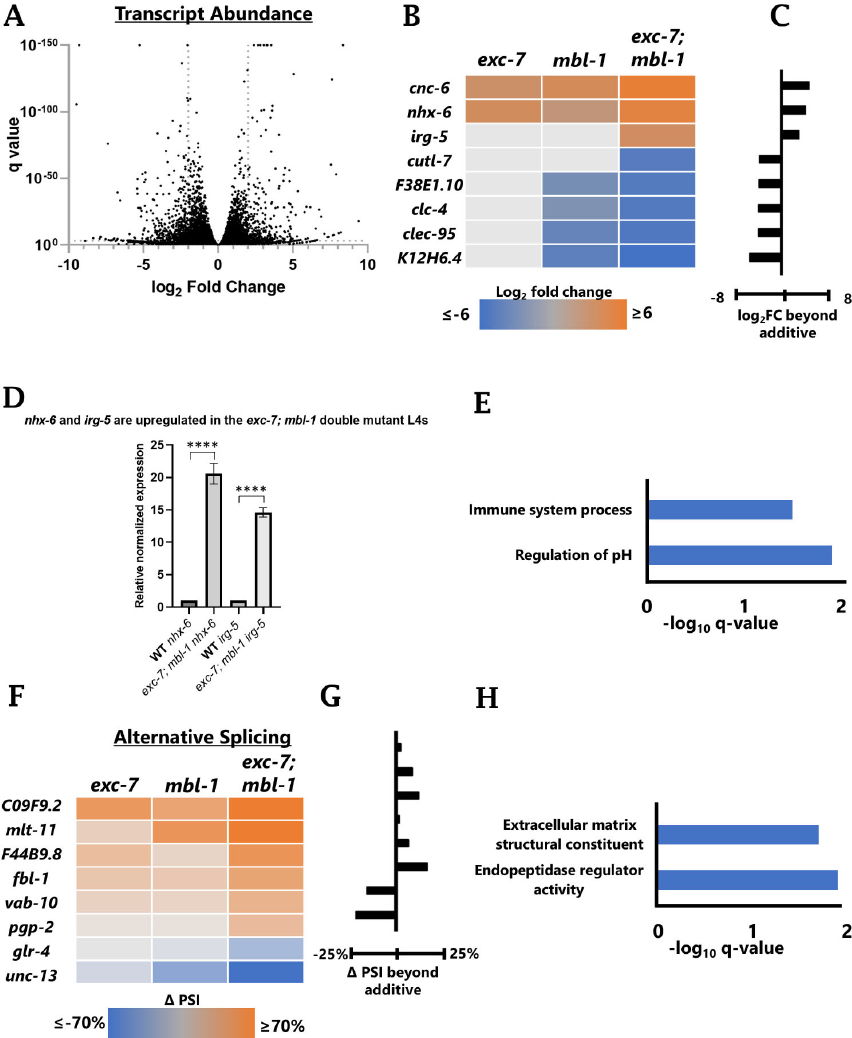
Unique gene dysregulation and splicing event changes in *exc-7; mbl-1* double mutants. (A) Volcano plot of gene expression changes between wild-type and *exc-7; mbl-1* double mutants. (B) Heat map of significantly dysregulated gene expression of single *exc-7*, *mbl-1*, and double mutant *exc-7; mbl-1* compared to wild type worms at the L4 stage. Blue indicates downregulation, orange upregulation in double mutant. Log 2-fold cutoff, P value 0.05 cutoff. (C) Gene expression changes that are greater than additive effects of *exc-7* and *mbl-1* single mutants. The excess gene expression change in the double mutant not accounted for by the sum of the single-mutant changes is plotted. (D) Relative normalized expression for *nhx-6* and i*rg-5* in *exc-7; mbl-1* double mutants normalized to wild type L4 worms. *exc-7; mbl-1* double mutants have increased expression of *nhx-6* and *irg-5* as predicted by our RNA seq analysis. T-test. *act-1*, *tba-1*, and *Y45F10D.4* used as housekeeping genes for ΔΔct normalization. (E) Gene Ontology analysis for genes with synthetically dysregulated expression levels. (F) Heat map of significantly changed alternative splicing events of single *exc-7*, *mbl-1*, and double mutant *exc-7; mbl-1* compared to wild type worms at the L4 stage. Blue indicates more exclusion, orange more inclusion in double mutant. (G) Splicing changes that are greater than additive effects of *exc-7* and *mbl-1* single mutants. The excess splicing change in double mutants not accounted for by the sum of the single-mutant changes is plotted. PSI = Percent Spliced in. (H) Gene Ontology analysis for genes with synthetically dysregulated splicing levels. For all panels, ns= P>0.05, *= P≤0.05, **= P≤0.01, ***= P≤0.001, ****=P≤0.0001.

We therefore filtered for changes in gene expression and alternative splicing that are more severe in the double mutant than in either single mutant. We further filtered for changes that are more severe in the double mutant than would be expected by additive effects of both *exc-7* and *mbl-1* single mutants. Using conservative cutoff thresholds (See Methods), we thus identified a small handful of genes dysregulated in a synthetic manner in *exc-7; mbl-1* double mutants. This is in contrast with hundreds of dysregulated genes in double mutants when compared directly with wild type. Filtering according to our conservative thresholds yields a total of eight synthetically regulated genes at the level of transcript abundance, as well as eight synthetically regulated alternative splicing events (Figure 3B-G).

We cross-validated some of the strongest dysregulated genes using RT-PCR for alternatively-spliced genes and qRT-PCR for genes regulated at the level transcript abundance (Figure 3D, S3). Gene ontology analysis of dysregulated genes identified different functional categories for genes dysregulated at the transcript level (immune and pH regulation) compared to those with dysregulated alternative splicing (extracellular matrix and endopeptidase regulator) (Figure 3E and 3H). In sum, our filtering strategy allowed us to substantially narrow our focus to genes dysregulated synthetically by *exc-7* and *mbl-1* to prioritize for follow-up functional analysis.

### Functional analysis of genes dysregulated in *exc-7; mbl-1* double mutants

Our stringent list of genes synthetically dysregulated in *exc-7; mbl-1* was sufficiently small for us to systematically test each gene for functional impact on lifespan using genetics (Fig 4). We used existing loss-of-function mutant alleles when available, generated over-expression constructs, and generated new CRISPR-mediated deletions for genes with no existing loss-of-function alleles (Supplemental table 1).

**Figure 4:**
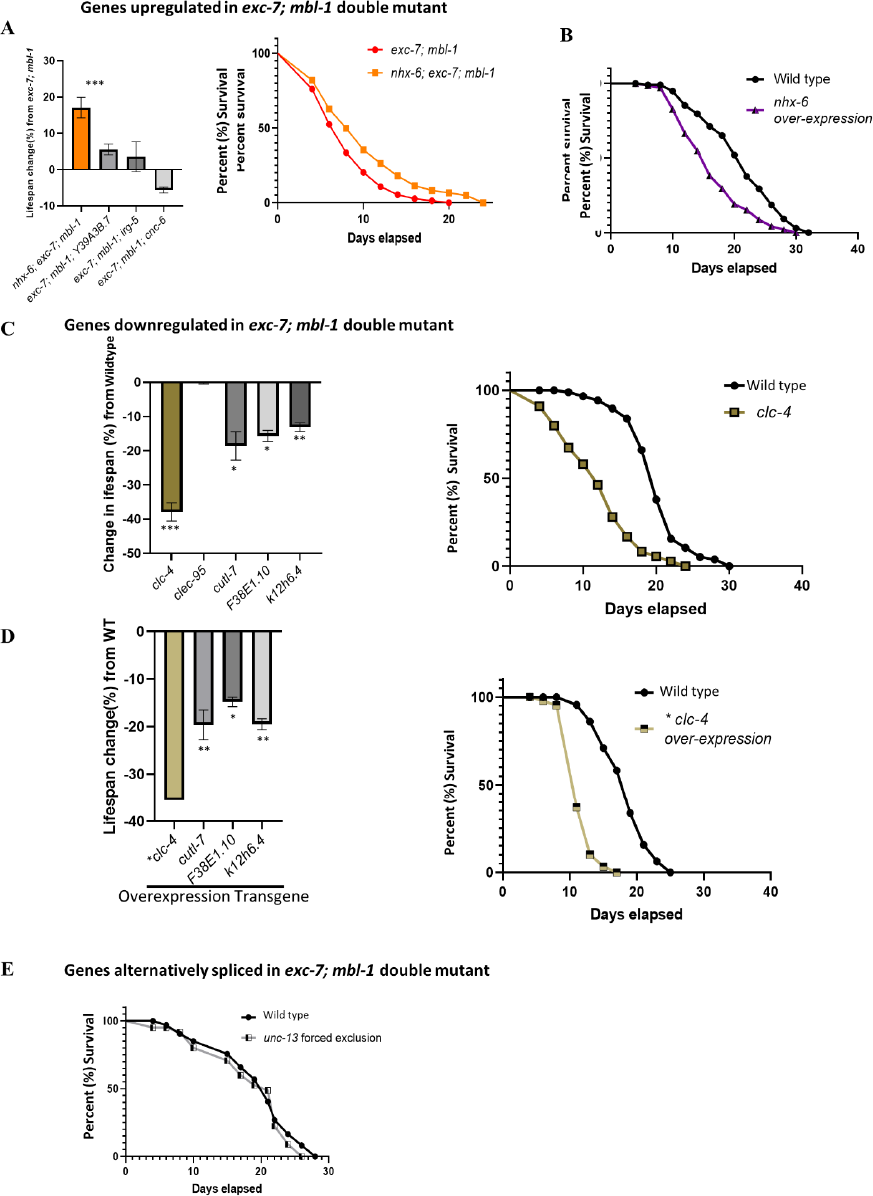
*nhx-6* is partially responsible for *exc-7; mbl-1* lifespan phenotype. (A) Triple mutant lifespan changes (%) from *exc-7; mbl-1* double mutant. Additional biological replicates in Figure S4 Three of the four triple mutants show no statistically-significant change in lifespan. Representative *nhx-6(ok609); exc-7; mbl-1* lifespan assay. The fourth triple mutant, *nhx-6; exc-7; mbl-1*, displays a significantly longer lifespan than an *exc-7; mbl-1* double mutant. P-value= *exc-7; mbl-1* vs *nhx-6(ok609); exc-7; mbl-1*\*\* (log-rank test). (B) Representative *nhx-6* O.E aging assay. Transgenic over-expression of *nhx-6* leads to a decrease in lifespan in wild-type animals. P-value= **** (log-rank test). (C) Single deletion mutant lifespan changes (%) from wild type. For genes with decreased expression in the *exc-7; mbl-1* double mutant, we predict that if under-expression of that gene contributes to the lifespan defect, then deleting that gene in a wild-type background would lead to decreased lifespan. None of the genes we tested meet these criteria. (One-way ANOVA). Representative *clc-4* lifespan assay. P-value**** (log-rank test). (D) Overexpression (O.E.) effects on lifespan - changes (%) from wild type. For genes with decreased expression in the *exc-7; mbl-1* double mutant, we predict over-expressing them in wild type background might lead to lifespan extension. None of the genes we tested meet these criteria (one-way ANOVA). Representative *clc-4* GFP fosmid lifespan assay. P-value**** (log-rank tes)t (E) Representative *unc-13* splicing lifespan assay. Dysregulated splicing of *unc-13* does not on its own contribute to the *exc-7; mbl-1* lifespan phenotype. Bonferroni P-values: wild type vs *unc-13* forced exclusion(ns) (log-rank test). For all panels, ns= P>0.05, *= P≤0.05, **= P≤0.01, ***= P≤0.001, ****=P≤0.0001.

For genes with increased expression in *exc-7; mbl-1* mutants, we predict that if overexpression of that gene contributes to the lifespan defect, then deleting it in the context of an *exc-7; mbl-1* double mutant would increase the lifespan of the double mutant. We also predict that overexpressing that gene in a wild-type background might decrease lifespan.

To test the first prediction, we generated a series of triple mutants that are mutant for *exc-7*, *mbl-1*, and one of the four upregulated genes (Figure 4A). Three of the four triple mutants show no statistically-significant change in lifespan (Figure 4A). But the fourth triple mutant, *nhx-6; exc-7; mbl-1*, displays a significantly longer lifespan than an *exc-7; mbl-1* double mutant (Figure 4A). This suggests that *nhx-6* overexpression contributes to the *exc-7; mbl-1* lifespan defect. We therefore tested the second prediction, that *nhx-6* overexpression would lead to a decrease in lifespan. Indeed, transgenic over-expression of *nhx-6* leads to a decrease in lifespan in wild-type animals (Figure 4B). Together this implicates *nhx-6* overexpression as an important contributor to the *exc-7; mbl-1* lifespan phenotype.

For genes with decreased expression in the *exc-7; mbl-1* double mutant, we predict that if under-expression of that gene contributes to the lifespan defect, then deleting that gene in a wild-type background would lead to decreased lifespan, while over-expressing it might lead to lifespan extension. None of the genes we tested meet these criteria (Figure 4C). While some of the single mutants do result in reduced lifespan, overexpressing these genes does not increase lifespan, but rather reduces lifespan (Figure 4D). Therefore, we did not prioritize any of the under-expressed genes for follow-up studies.

For genes with dysregulated splicing in the *exc-7; mbl-1* double mutant, we identified one particularly strong example of synthetic dysregulation in the gene *unc-13*. This alternatively-spliced exon is mostly included in wild type or in single mutants, but is largely skipped in the double mutant (Fig 3F). If this dysregulated splicing contributes to the lifespan defect of *exc-7; mbl-1*, then we would predict that disrupting *unc-13* splicing should result in decreased lifespan. To test this prediction, we used CRISPR/Cas9 to precisely excise the alternative exon from the genome, forcing it to be skipped. This manipulation does not result in a change in lifespan (Figure 4E), indicating that dysregulated splicing of *unc-13* does not on its own contribute to the *exc-7; mbl-1* lifespan phenotype.

Taken together, our systematic functional analysis of genes synthetically dysregulated in *exc-7; mbl-1* reveals a single target, *nhx-6*, as a promising candidate to partially explain the *exc-7; mbl-1* lifespan deficit. *nhx-6* is highly overexpressed in *exc-7; mbl-1* double mutants, deletion of *nhx-6* partially rescues the lifespan defect, and overexpression of *nhx-6* causes lifespan defects in a wild-type background.

### *nhx-6* expression affects lifespan and intestinal healthspan

To further characterize the role of *nhx-6* in mediating lifespan, we first validated that the increased lifespan observed in an *exc-7; mbl-1* double mutant is due to the on-target *nhx-6(ok609)* mutation and not due to a background mutation in the strain. To do this we used CRISPR/Cas9 to generate a new null allele, deleting the entire *nhx-6* gene. As was the case with the original *nhx-6* allele, our new deletion also significantly increases lifespan in an *exc-7; mbl-1* mutant, but not in a wild-type background (Figure 5A-B). In fact, in a wild-type background, loss of *nhx-6* causes a modest decrease in lifespan (Figure 5A-B). This further supports the role of aberrant *nhx-6* overexpression as a contributor to the *exc-7; mbl-1* lifespan phenotype, and indicates that precise regulation of *nhx-6* levels are important for lifespan.

*nhx-6* has previously been shown to be expressed in the intestine, where it functions as a sodium/proton exchanger and likely controls pH and/or sodium homeostasis, although no clear phenotype or functional role has yet been identified (21). Due to its established role in the intestine, we first tested whether *nhx-6* contributes to the intestine-related phenotypes observed in the *exc-7; mbl-1* double mutants. We found that intestinal permeability is affected by *nhx-6* levels: overexpression of *nhx-6* results in an increase in intestinal permeability, similar to the *exc-7; mbl-1* double mutant (Figure 5C). Moreover, deleting *nhx-6* partially rescues the permeability phenotype of the *exc-7; mbl-1* double mutant (Figure 5C).

**Figure 5:**
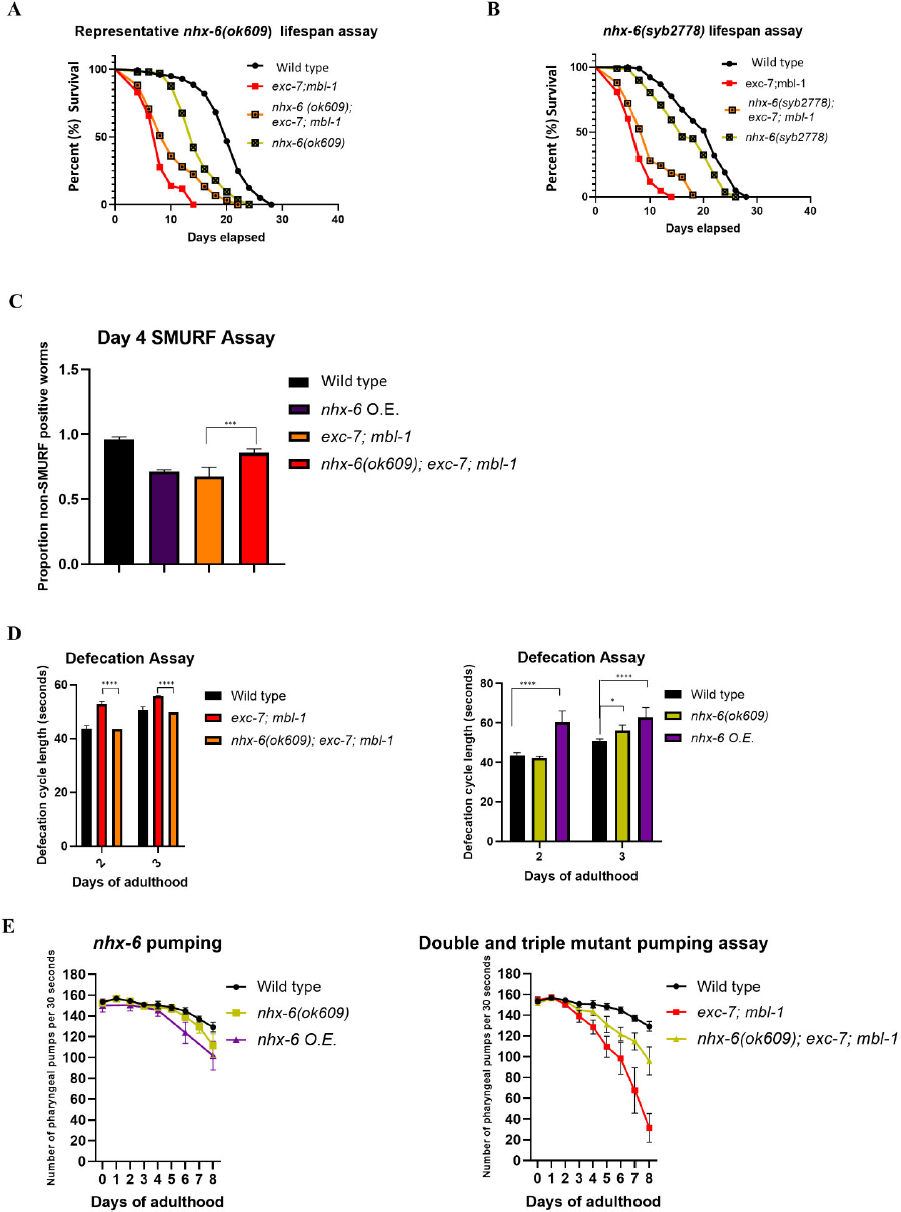
*nhx-6* expression levels affect lifespan and healthspan phenotypes. (A) Representative *nhx-6(ok609)* lifespan assay. *nhx-6* deletion significantly increases lifespan in an *exc-7; mbl-1* mutant, but not in a wild-type background. Additional biological replicates in Figure S5. Bonferroni P-values: wild type vs *nhx-6(ok609)*\*\*\*\*, *exc-7; mbl-1* vs *nhx-6(ok609); exc-7; mbl-1*\*\*, (log-rank test). (B) Representative *nhx-6(syb2778)* lifespan assay. Our new deletion also significantly increases lifespan in an *exc-7; mbl-1* mutant, but not in a wild-type background Bonferroni P-values: wild type vs *nhx-6(syb2778)*\*\*, *exc-7; mbl-1* vs *nhx-6(syb2778); exc-7; mbl-1*\*, (log-rank test). (C) SMURF/ intestinal permeability assay Day 4. Over expressing *nhx-6* increases intestinal permeability compared to wild type worms. *nhx-6* deletion mutants have a similar SMURF rate to wild type worms. P-value= *exc-7; mbl-1* vs *nhx-6(ok609); exc-7; mbl-1*\*\*\*. n= *exc-7; mbl-1* n= 88, 130, 50, *nhx-6(ok609); exc-7; mbl-1* n= 90, 100, 55. (Chi-squared test). (D) Defecation assay. Deletion of *nhx-6* in a wild-type background does not affect the defecation cycle, but deleting *nhx-6* in an *exc-7; mbl-1* background partially rescues the double mutant phenotype. P-values: day 2-3 *exc-7; mbl-1* vs *nhx-6; exc-7; mbl-1,* (one-way ANOVA). (E) *nhx-6* pumping assays. Deleting *nhx-6* in a wild-type background results in no change in pumping activity, but deleting *nhx-6* in an *exc-7; mbl-1* background partially rescues the progressive pumping defects. P-values: L4-day 2: all (ns), day 3: wild type vs *nhx-6(ok609)*(ns), *exc-7; mbl-1* vs *nhx-6(ok609); exc-7; mbl-1*(ns), day 4: wild type vs *nhx-6(ok609)*(ns), day 5: wild type vs *nhx-6(ok609)*(ns), day 6: wild type vs *nhx-6(ok609*)(ns), day 7: wild type vs *nhx-6(ok609)*(ns), day 8: wild type vs *nhx-6(ok609)*(ns), (one-way ANOVA). For all panels, ns= P>0.05, *= P≤0.05, **= P≤0.01, ***= P≤0.001, ****=P≤0.0001.

As another test for intestinal function, we assayed the defecatory motor program in double mutants. *C. elegans* defecation occurs in a stereotyped periodic cycle of roughly 50 seconds per cycle, and is controlled by the intestine together with two neurons and small number of muscles (22). We found that *exc-7; mbl-1* mutants have longer defecation cycles, as do worms overexpressing *nhx-6* (Figure 5D). Deletion of *nhx-6* in a wild-type background does not affect the defecation cycle, but deleting *nhx-6* in an *exc-7; mbl-1* background partially rescues the double mutant phenotype (Figure 5D), indicating that *nhx-6* overexpression is partially responsible for the progressive intestinal defects observed in *exc-7; mbl-1* mutants.

*exc-7; mbl-1* mutants exhibit progressive defects in pumping in the pharynx, which is responsible for capturing food and transporting it to the intestine. As such, pharyngeal activity defects could in principle originate from the intestine, or from the cells of the pharynx itself. We found that *nhx-6* overexpression causes modest pumping defects that increase with age (Figure 5E). Deleting *nhx-6* in a wild-type background results in no change in pumping activity, but deleting *nhx-6* in an *exc-7; mbl-1* background partially rescues the progressive pumping defects (Figure 5E). Together, these results indicate that *nhx-6* expression has an effect on lifespan, as well as aspects of intestinal healthspan, and suggest that intestinal defects in *exc-7; mbl-1* mutants are in part due to *nhx-6* overexpression.

### *nhx-6* expression affects specific systemic healthspan phenotypes

We next tested whether *nhx-6* overexpression might contribute systemically to non-intestinal phenotypes in *exc-7; mbl-1* double mutants. We tested various phenotypes observed in the *exc-7; mbl-1* double mutants, and asked whether *nhx-6* mutation would rescue any of the phenotypes. For example, we observed that *exc-7; mbl-1* mutants exhibit a substantial reduction in fertility, but *nhx-6* deletion does not rescue this phenotype (Figure 6A). We also noticed that young adult *exc-7; mbl-1* worms are smaller than wild-type worms, and that *nhx-6* mutation results in a small but significant rescue of the size phenotype in *exc-7; mbl-1* double mutants (Figure 6B).

**Figure 6:**
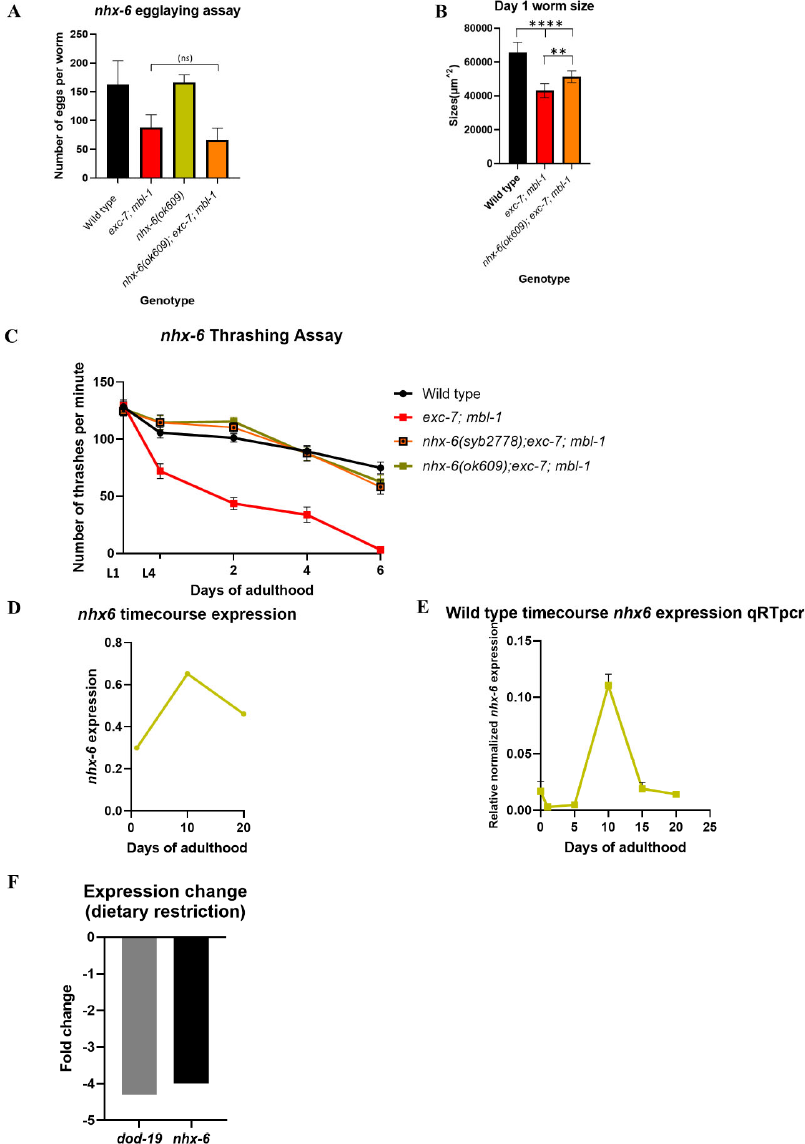
*nhx-6* partially rescues lifespan and healthspan defects in *exc-7; mbl-1* mutants. (A) nhx-6 egglaying assay. *exc-7; mbl-1* mutants exhibit a substantial reduction in progeny production, but *nhx-6* deletion does not rescue this phenotype. P-values: *exc-7; mbl-1* vs *nhx-6(ok609); exc-7; mbl-1*(ns), (one-way ANOVA). (B) D1 sizes: Young adult *exc-7; mbl-1* worms are smaller than wild-type worms, and that *nhx-6* mutation results in a small but significant rescue of the size phenotype in *exc-7; mbl-1* double mutants (one-way ANOVA). (C) *nhx-6* thrashing assay. Number of thrashes per minute. *nhx-6* mutation results in a nearly complete rescue of locomotion defects throughout the animal’s lifespan. P-values: L1: all (ns), L4-D6: *exc-7; mbl-1* vs *nhx-6(syb2778); exc-7; mbl-1*\*\*\*\*, *exc-7; mbl-1* vs *nhx-6(ok609); exc-7; mbl-1*\*\*\*\*, (one-way ANOVA). (D) Published RNAseq data has shown increased *nhx-6* expression with age. We examined previously published RNA Seq timecourse data (23) and found that *nhx-6* expression increases with age, peaking around day 10 of adulthood, which is roughly the midpoint of adult lifespan. (E) Time course qRT PCR conducted on wild type worms similarly showed an increase in *nhx-6* with age. We performed qRT-PCR on wild type worms at additional timepoints and confirmed a striking increase in *nhx-6* levels, particularly at day 10 of adulthood. (F) RNA Seq data showing *nhx-6* expression decreases under dietary restriction conditions. We examined RNA Seq data performed under dietary restriction, which extends lifespan in *C. elegans* (24,25), and found that *nhx-6* expression decreases under dietary restriction. *dod-19* is shown in comparison as a positive control gene that has previously been shown to be sensitive to dietary restriction and a downstream target of DAF-16/FOXO. For all panels, ns= P>0.05, *= P≤0.05, **= P≤0.01, ***= P≤0.001, ****=P≤0.0001.

Finally, we tested locomotory ability and found that *exc-7; mbl-1* double mutants have strong and progressive locomotion defects (Figure 6C). Strikingly, *nhx-6* mutation results in a nearly complete rescue of these defects throughout the animal’s lifespan (Figure 6C). Together, these results indicate that many, but not all, *exc-7; mbl-1* phenotypic defects are relieved by *nhx-6* mutation. This suggests that overexpression of *nhx-6* comprises a major part of the *exc-7; mbl-1* lifespan and healthspan phenotype.

### *nhx-6* in aging

Having demonstrated that *nhx-6* mutants partially rescue the lifespan and healthspan phenotypes of *exc-7; mbl-1* double mutants (Figure 5-6), we next asked whether *nhx-6* expression might be associated with aging in wild type worms. We first examined previously published RNA Seq time course data (23) and found that *nhx-6* expression increases with age, peaking around day 10 of adulthood, which is roughly the midpoint of adult lifespan (Figure 6D). We further confirmed and extended this observation by performing qRT-PCR on wild type worms at additional timepoints (Figure 6E) and confirmed a striking increase in *nhx-6* levels, particularly at day 10 of adulthood. Finally, we examined RNA Seq data performed under dietary restriction, which extends lifespan in *C. elegans* (24,25), and found that *nhx-6* expression decreases under dietary restriction (Figure 6F). Together these results indicate that not only does *nhx-6* partially rescue the lifespan and a number of healthspan phenotypes seen in the *exc-7; mbl-1* double mutant, but that *nhx-6* expression is also associated with aging processes in wild-type worms.

## DISCUSSION

We demonstrate that a pair of RBPs are required for lifespan and healthspan in *C. elegans*. These findings are in accordance with emerging evidence that RNA regulation by RBPs is critical for lifespan, in addition to the well-established roles for transcription factors in modulating lifespan (4–7). Recent work in *C. elegans* has led to the hypothesis that the lifespan phenotypes observed in RBP mutants is caused by alternative splicing dysregulation (4,5). We find that *exc-7; mbl-1* mutants result in synthetically dysregulated alternative splicing, but have not as yet identified a causative link between this dysregulated splicing and the double mutant phenotype. In contrast, we do provide evidence for a link between RNA abundance of *nhx-6* and the double mutant phenotype. This highlights the fact that RBPs have myriad regulatory activities, and their functional outputs may result from a combination of different types of regulatory activities, including some not addressed by our study, *e.g.* translational control or RNA localization. In the case of *nhx-6* regulation, the precise mechanism of regulation (direct vs. indirect binding, co-transcriptional vs. post-transcriptional, etc.) await further investigation.

The strategy that we deployed for identifying uniquely dysregulated genes in double mutants that are not explained by the constituent single mutants could be extended to other similar use cases. Synthetic genetic interaction between gene regulatory factors is a widely-observed phenomenon (19,26–28), and the ability to distill transcriptomic networks down to a handful of synthetically-regulated candidates is a promising approach for identifying functionally relevant regulatory targets.

In the present study, our analysis identified a single target, *nhx-6*, whose overexpression is partially responsible for the *exc-7; mbl-1* lifespan phenotype. How might this overexpression cause the observed lifespan and healthspan defects? *nhx-6* encodes an ion antiporter that is predicted to perform Na+-H+ ion exchange across the plasma membrane. The function of *nhx-6* remains uncharacterized, except for the observation that it is expressed in the intestine, where it is localized basolaterally at the posterior and anterior ends of the intestine, but localized apically in the middle of the intestine (21).

As an intestinal ion transporter, the observed *nhx-*6-mediated defects in intestinal phenotypes may be direct effects of altered ion homeostasis in the intestine. For example, intestinal pH dynamics play a critical role in regulating the defecation cycle (15,29,30), and defects in intestinal permeability could be a direct consequence of prolonged lack of ion homeostasis in the intestine. Indeed, loss of another intestinal ion transporter, *nhx-2*, has been shown to cause intestinal pH imbalance and decreased lifespan (31). Loss of intestinal permeability is correlated with lifespan in a range of diverse organisms, and is an accurate predictor of mortality in a number of animal models of aging (15,29,30). We therefore speculate that intestinal barrier dysfunction might represent a primary driver of *nhx-6-*related loss of healthspan and lifespan, while additional phenotypes such as pharyngeal pumping and fertility might result from downstream consequences of intestinal barrier dysfunction. Future work will address this question and further delineate the links between combinatorial RBP function, *nhx-6* overexpression, and *C. elegans* lifespan.

## METHODS

### Strains used/generated

See table S1 for list of strains used and generated. Some strains were provided by the C.G.C., which is funded by NIH Office of Research Infrastructure Programs (P40 OD010440). Strains with PHX designation were generated by SunyBiotech.

### Primers used

See tableS2.

### Lifespan assay

Staged L4 worms (n= 100 worms per genotype per assay) grown at 20°C were picked to NGM + FUDR (50 µg/mL) plates seeded with OP50 bacteria and grown at 20°C. Worms were considered dead when they ceased all spontaneous movement and no longer responded to touch from a platinum wire. Number of live, dead, and censored animals were recorded every two days. Statistical significance was assessed by performing a log-rank test (32). Assays were performed in triplicate.

### Pharyngeal pumping assay

Pharyngeal pumping assays were adapted from (33). Briefly, staged L4 worms grown at 20°C were picked to NGM + FUDR (50 µg/mL) plates seeded with OP50 bacteria and grown at 20°C. At each timepoint the pharyngeal pumping rate was manually recorded for 30 seconds. (t-test). n= 5 worms per genotype per timepoint. Assays conducted in triplicate.

### SMURF/ Intestinal permeability assay

Adapted from (34). Briefly, staged L4 worms grown at 20°C were plated on NGM + FuDR (50 µg/mL) plates seeded with OP50 bacteria and grown at 20°C. At each timepoint 50+ Worms were picked off plates and suspended in a liquid suspension (1:1) of M9 (KH2PO4 -– Na2HPO4 -– NaCl -– NH4Cl) buffer mixed with blue food dye (Erioglaucine disodium salt, Sigma-Aldrich, 5.0% wt/vol (5g/100mL) in water (stock) for 3 h.

Animals were then washed three times with M9 buffer allowing for the worms to gravity settle between washed. 20 μL of the pellet was placed on a microscope slide for viewing. 10 μL Sodium azide (20 mM) was added to paralyze the worms. The worms were visually analyzed for the presence and/or the absence of leaked blue dye outside the intestinal lumen and throughout the body cavity using microscope imaging. The percentage of “non-smurf” worms was calculated per genotype at each timepoint.

### Imaging

Fluorescent Images taken at 10X using a Zeiss Axio Imager.Z1 AX10 microscope and Zeiss ZEN2.5 (blue edition) software. Images were processed using ImageJ 1.54d (NIH, USA. http://imagej.org). White-light images for SMURF assay taken with AMScope Microscope digital camera (MU300-HS) and AmLite, Copyright © 2014 – 2018.

### Egg laying assay

Five L4 worms of each genotype were placed on 3 seeded NGM plate and incubated at 20°C. The following day, and every subsequent day until egg production stopped, the 5 adults were transferred to new seeded plates. The number of progeny on each plate was counted and divided by the total number of adults to determine average egg production per day per worm. Total brood size was quantified as the sum total of eggs produced over lifespan per worm. Assays conducted in triplicate.

### Thrashing/swimming assay

#### Worm lab assays (wave initiation rate)

Staged L4 worms grown at 20°C were picked to NGM + FUDR (50 µg/mL) plates seeded with OP50 bacteria and grown at 20°C. At each timepoint ∼30 worms per genotype were picked over to non-seeded NGM plates. These worms were allowed to crawl around for 5+ minutes to remove bacteria. Worms were picked in groups of 5-10 worms into a 20 μL drop of M9 buffer. This allowed for multiple tracks to be recorded within one capture but insured that the worms were spread-out enough in order to not overlap and allowing for clean captures. Three 1-minute recordings were made per genotype per timepoint. Videos were then processed according to WormLab suggestions 10 clean tracks per genotype per timepoint were used to calculate the average thrashing rate (wave initiation rate). n= 10 tracks per genotype per timepoint. Assays conducted at least in triplicate. Worms maintained on FuDR (50 mg/mL) plates at 20°^C^. Assays conducted at room temperature using WormLab. (WormLab 2022. (MBF Bioscience LLC, Williston, VT USA). WormLab®(2022.1.1)64-bit.

### Height and width measurements

Images were obtained on a Zeiss Axio Imager.Z1 AX10 microscope and Zeiss ZEN2.5 (blue edition) software (©<u>Carl Zeiss Microscopy</u> GmBh, 2018) was used to obtain measurements. Heights were measured along center line from nose to tail. Width measurements were taken at vulva point.

### RNAseq analysis

Total RNA was extracted from L4 worms using Tri reagent according to manufacturer’s protocol (Sigma Aldrich). Three biological replicates were extracted per genotype. mRNA was purified from each sample using NEBNext® Poly(A) mRNA Magnetic Isolation Module, and cDNA libraries were prepared using NEBNext® Ultra™ II RNA Library Prep Kit for Illumina, following kit protocols. Libraries were sequenced on Illumina HiSeq 2000, paired-end 150 bp reads. Raw fastq files are available at the NCBI SRA (PRJNA862903).

Total RNA was extracted from L4 stage worms using Tri reagent (Sigma Aldrich) as recommended by the manufacturer. A total of three biological replicates were collected for each sample. PolyA+ transcripts were converted to cDNA libraries using the TruSeq RNA kit (Illumina). Libraries were sequenced on Illumina HiSeq 2000, paired-end 150 bp reads, then mapped to the worm genome (versionWBcel235) using STAR(35). Gene-specific counts were tabulated for each sample using HTSeq (36) and statistically-significant differentially expressed transcripts were identified with DESeq2(37). The Junction Usage Model (2.0.2) was used to identify differentially spliced isoforms in experimental samples compared to wild type controls and quantify their expression levels by computing the ΔPSI (difference of Percent Spliced Isoform)(38).

Identification of uniquely dysregulated genes was performed in two filtering steps, which we designed to be strict/conservative thresholds. For the first filtering step, genes dysregulated at the level of RNA abundance were required to pass statistical and effect-size thresholds in comparisons between *exc-7; mbl-1* double mutants and wild type animals. For gene expression, this was q-value <0.01 and |log_2_FC| >1 as determined by DESeq2. For alternative splicing this was q-value <0.01 and |ΔPSI| > 10% as determined by JUM. The second filtering step was a requirement that the magnitude of dysregulation was (a) greater in the double mutant than either of the single mutants and (b) greater in the double mutant than in the sum of both single mutants.

Custom scripts used for this pipeline can be found at <u>github.com/TheNorrisLab/Double_Mutant_Gene_Expression_Analysis</u>. For gene expression, we required the difference to be > log_2_FC=1, and for alternative splicing, ΔPSI > 10%.

### Gene ontology

Performed with WormBase’s Enrichment Analysis tool (39,40).

### qRT PCR

Worms were staged and maintained at 20°C. At L4 RNA was extracted using Tri-reagent and Direct-zol RNA MiniPrep Kit (Zymo research, Genesee Scientific) as per kit recommendations. cDNA made using Verso cDNA Synthesis Kit (ThermoFisher) as per kit recommendations. 500ng RNA per 20 uL cDNA synthesis reaction. act-1, tba-1, and Y45F10D.4 were used as housekeeping genes. 1uL cDNA used per reaction with (ThermoFisher) PowerUp SYBR Green Master Mix.

### Defecation assay

Adapted from (41). Briefly, Staged L4 worms (n= 100 worms per genotype per assay) grown at 20°C were picked to NGM + FUDR (50 µg/mL) plates seeded with OP50 bacteria and grown at 20°C. At each time point individual worms were observed 5 minutes as the duration between expulsions were recorded. n=10 per genotype per timepoint. Assays conducted in triplicate.

### Statistical analysis

Lifespan Assays:

Survival plots, Restricted means (days), and Bonferroni P-values calculated via Log-Rank Test using Oasis2 (32).

Pharyngeal Pumping Assays:

t-test and one-way ANOVA. n= 5 worms per genotype per timepoint. Assays conducted in triplicate. SMURF/ intestinal permeability assay:

Welch’s t test. n= 50+ worms per genotype per time point. Assays conducted in triplicate. Day 4. Chi-squared test. n= *exc-7; mbl-1* n= 88, 130, 50, *nhx-6(ok609); exc-7; mbl-1* n= 90, 100, 55. Assays conducted in triplicate.

Thrashing Assay:

Welch’s t test. n= 10 worms per genotype per timepoint. Assays conducted at least in triplicate.

Relative normalized expression.:

T-test. Triplicates were performed within 3 technical replicates with 3 biological replicates. *act-1*, *tba-1*, and *Y45F10D.4* used as housekeeping genes for ΔΔct normalization (see tableS2 for primer sequences).

Defecation assay:

one-way ANOVA. Assays conducted in triplicate.

Egg laying assay:

one-way ANOVA. Assays conducted in triplicate.

Worm size:

one-way ANOVA. Assays conducted in triplicate.

## ACKNOWLEDGEMENTS

Thank you to members of the Norris Lab and to Megan Norris for critical reading of the manuscript and for helpful suggestions.

---

**Supplemental Table 1.**
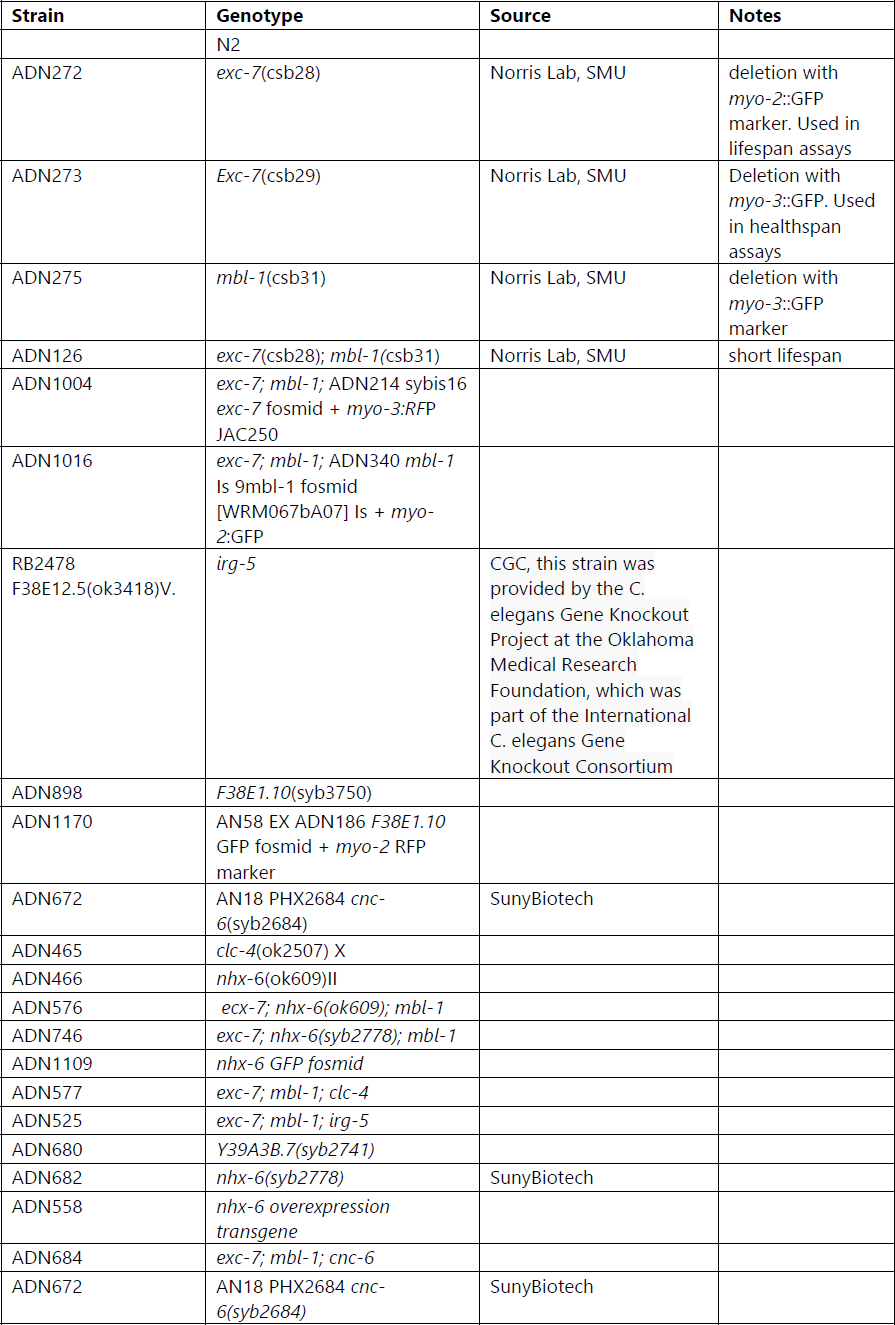
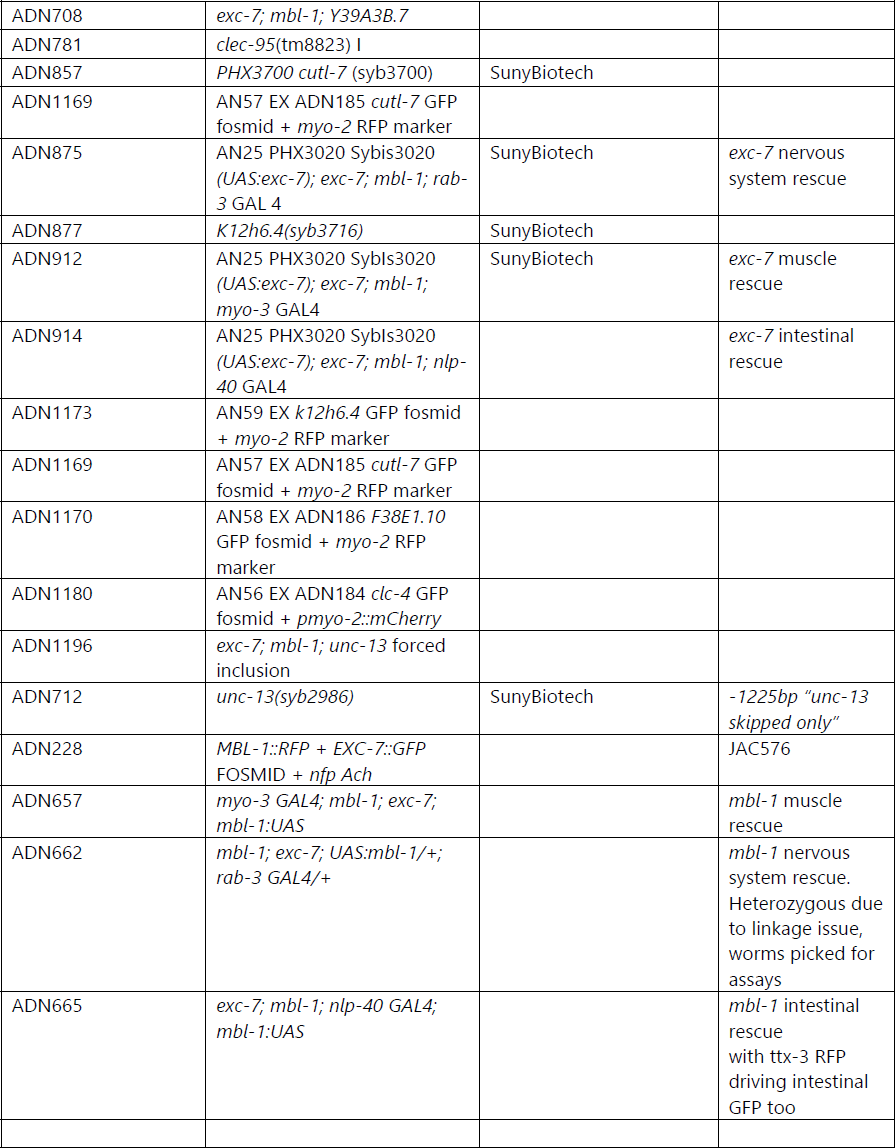
Strains used.

**Supplemental Table 2.**
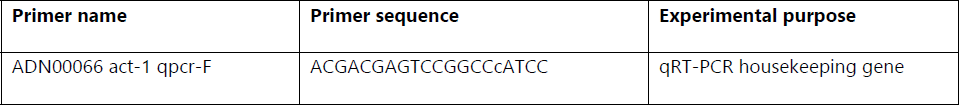
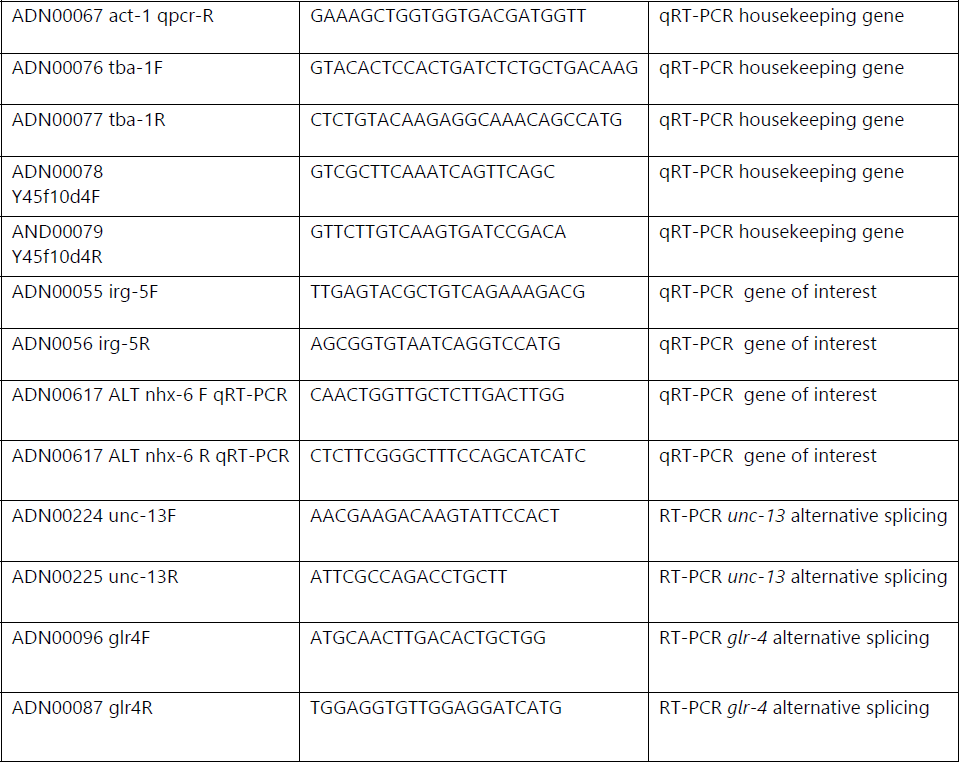
Primers used.

## Supplemental figures

**Supplemental figure 1.**
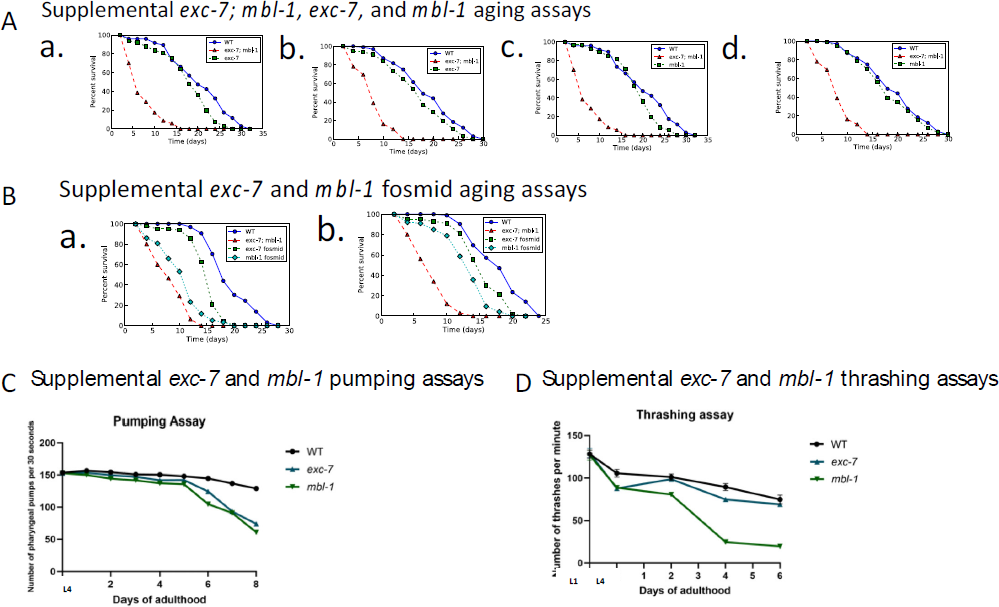
A. Supplemental *exc-7; mbl-1, exc-7, mbl-1* lifespan assays and *exc-7* and *mbl-1* fosmid lifespan assays. *exc-7; mbl-1, exc-7, mbl-1* lifespan assays: Bonferroni P-values: a. wild type vs *exc-7; mbl-1*= 0.0e+00, wild type vs *exc-7*= 0.0123, *exc-7; mbl-1* vs *exc-7*= 0.0e+00. b. wild type vs *exc-7; mbl-1*= 0.0e+00, wild type vs *exc-7*= 0.0681, *exc-7; mbl-1* vs *exc-7*= 0.0e+00. c. wild type vs *exc-7; mbl-1*= 0. 0e+00, wild type vs *mbl-1*= 0.0499, *exc-7; mbl-1* vs *mbl-1*= 0.0e+00. d. wild type vs *exc-7; mbl-1*= 0.0e+00, wild type vs *mbl-1*= 0.5897, *exc-7; mbl-1* vs *mbl-1*= 0.0e+00. B. *exc-7* and *mbl-1* fosmid lifespan assays: Bonferroni P-values: a. wild type vs *exc-7; mbl-1*= 0.0e+00, wild type vs *exc-7* fosmid= 0.0e+00, wild type vs *mbl-1* fosmid= 0.0e+00, *exc-7; mbl-1* vs *mbl-1* fosmid= 0.0001. b. wild type vs *exc-7; mbl-1*= 0.0e+00, wild type vs *exc-7* fosmid= 0.0e+00, wild type vs *mbl-1* fosmid= 0.0e+00. WT (wild type). C. *exc-7* and *mbl-1* pumping assays. P-values: L4: all= ns, D1: N2 vs *exc-7*= ns, N2 vs *mbl-1*= ****, D2: N2 vs *exc-7*= ns, N2 vs *mbl-1*= ***, D3-4: N2 vs *exc-7*= ns, N2 vs *mbl-1*= **, D5: N2 vs *exc-7*= ns, N2 vs *mbl-1*= *, D6: N2 vs *exc-7*= ns, N2 vs *mbl-1*= ****, D7: all ****, D8: all ****. One-way ANOVA. D. *exc-7* and *mbl-1* thrashing (wave initiation rate) assays (thrashes per minute). P-values: L1= all ns, L4=N2 vs *exc-7*= ****, N2 vs *mbl-1*= ****, D2: N2 vs *exc-7*= ns, N2 vs *mbl-1*= ****, D4: N2 vs *exc-7*= **, N2 vs *mbl-1*= ****, D6: N2 vs *exc-7*= ns, N2 vs *mbl-1*= ****. One-way ANOVA. For all panels, ns= P>0.05, *= P≤0.05, **= P≤0.01, ***= P≤0.001, ****=P≤0.0001.

**Supplemental figure 2.**
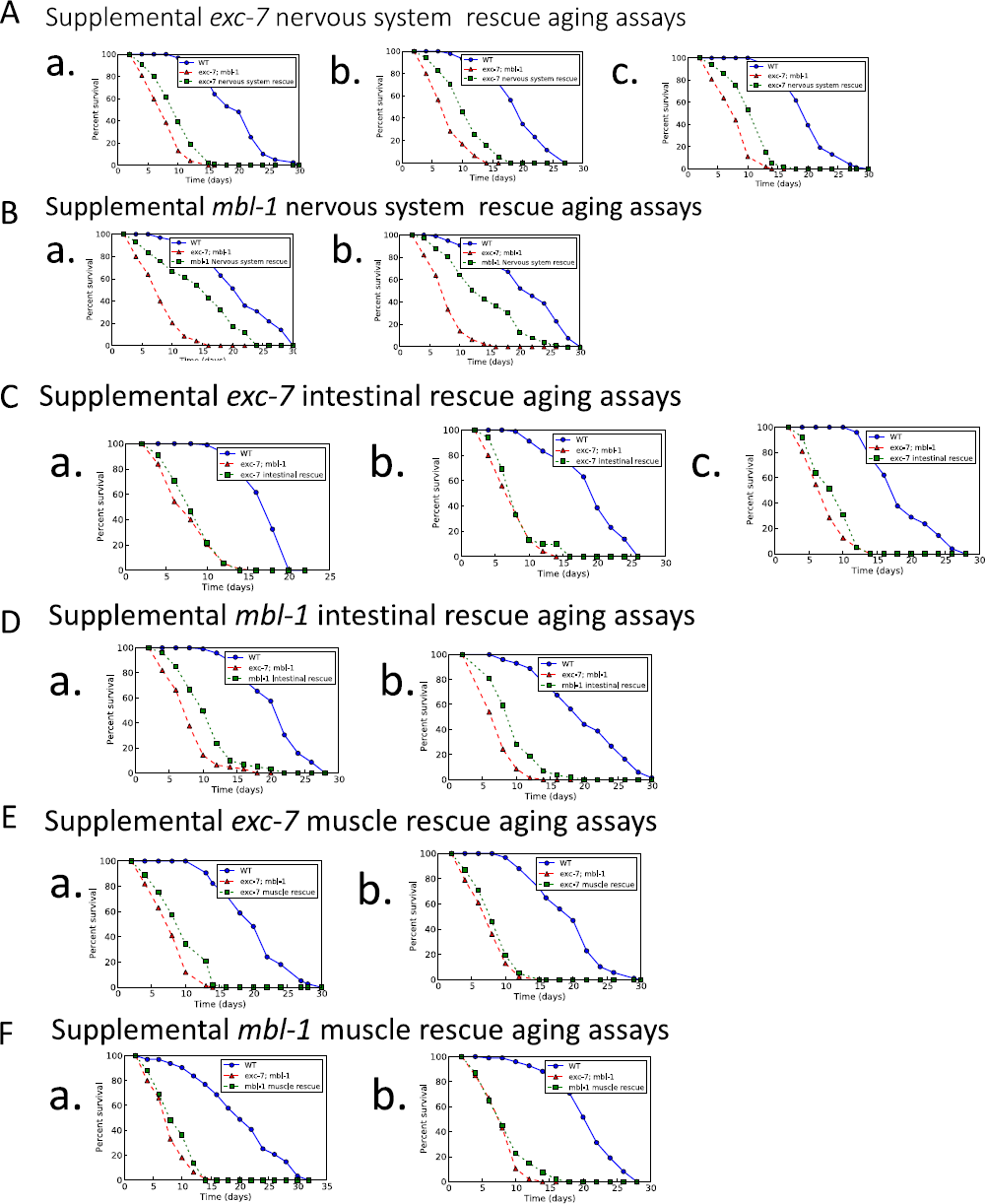
A. Tissue specific rescue aging assays. Supplemental *exc-7* nervous system rescue lifespan assays. Bonferroni P-values: a. wild type vs *exc-7; mbl-1*= 0.0e+00, wild type vs *exc-7* nervous system rescue= 0.0e+00, *exc-7; mbl-1* vs *exc-7* nervous system rescue= 1.2e-05. b. wild type vs *exc-7; mbl-1*= 0.0e+00, wild type vs *exc-7* nervous system rescue= 0.0e+00, *exc-7; mbl-1* vs *exc-7* nervous system rescue= 1.4e-07. c. wild type vs *exc-7; mbl-1*= 0.0e+00, wild type vs *exc-7* nervous system rescue= 0.0e+00, *exc-7; mbl-1* vs *exc-7* nervous system rescue= 0.0e+00. **B. Supplemental *mbl-1* nervous system rescue lifespan assays.** Bonferroni P-values: a. wild type vs *exc-7; mbl-1*= 0.0e+00, wild type vs *mbl-1* nervous system rescue= 0.0e+00, *exc-7; mbl-1* vs *mbl-1* nervous system rescue= 0.0e+00. b. wild type vs *exc-7; mbl-1*= 0.0e+00, N2 vs *mbl-1* nervous system rescue= 0.0e+00, *exc-7; mbl-1* vs *mbl-1* nervous system rescue= 0.0e+00. **C. Supplemental *exc-7* intestinal rescue lifespan assays.** Bonferroni P-values: a. wild type vs *exc-7; mbl-1*= 0.0e+00, wild type vs *exc-7* intestinal rescue= 0.0e+00, *exc-7; mbl-1* vs *exc-7* intestinal rescue= 0.6197. b. wild type vs *exc-7; mbl-1*= 0.0e+00, wild type vs *exc-7* intestinal rescue= 0.0e+00, *exc-7; mbl-1* vs *exc-7* intestinal rescue= 0.2143. c. wild type vs *exc-7; mbl-1*= 0, wild type vs *exc-7* intestinal rescue= 0, *exc-7; mbl-1* vs *exc-7* intestinal rescue= 0.1670**. D. Supplemental *mbl-1* intestinal rescue lifespan assays.** Bonferroni P-values: a. wild type vs *exc-7; mbl-1*= 0.0e+00, wild type vs *mbl-1* intestinal rescue= 0.0e+00, *exc-7; mbl-1* vs *mbl-1* intestinal rescue= 1.4e-06. b. wild type vs *exc-7; mbl-1*= 0.0e+00, wild type vs *mbl-1* intestinal rescue= 0.0e+00, *exc-7; mbl-1* vs *mbl-1* intestinal rescue= 1.4e-07**. E. Supplemental *exc-7* muscle rescue lifespan assays.** Bonferroni P-values: a. wild type vs *exc-7; mbl-1*= 0.0e+00, wild type vs *exc-7* muscle rescue= 0.0e+00, *exc-7; mbl-1* vs *exc-7* muscle rescue= 0.0001. b. wild type vs *exc-7; mbl-1*= 0.0e+00, wild type vs *exc-7* muscle rescue= 0.0e+00, *exc-7; mbl-1* vs *exc-7* muscle rescue= 0.1481**. F. Supplemental *mbl-1* muscle rescue lifespan assays.** Bonferroni P-values: a. wild type vs *exc-7; mbl-1*= 0.0e+00, wild type vs *mbl-1* muscle rescue= 0.0e+00, *exc-7; mbl-1* vs *mbl-1* muscle rescue= 0.0475. b. wild type vs *exc-7; mbl-1*= 0.0e+00, wild type vs *mbl-1* muscle rescue= 0.0e+00, *exc-7; mbl-1* vs *mbl-1* muscle rescue= 0.1320.

**Supplemental figure 3.**
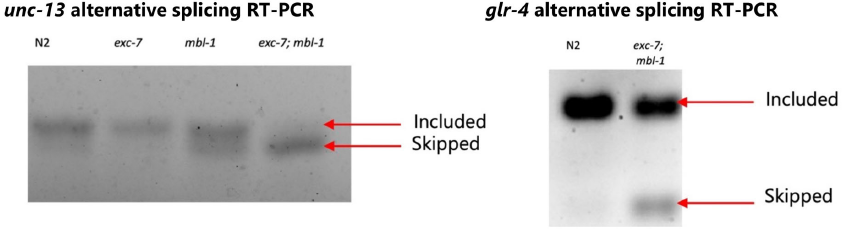
RT-PCR splicing event confirmation gels. *unc-13* splicing gel. Included= 82bp, skipped= 73bp. RNA-seq analysis suggests more skipped in *exc-7; mbl-1* which is confirmed in the RT-PCR gel. *glr-4* splicing gel. Included= 108bp, skipped= 86bp. RNA-seq analysis suggested more skipped in *exc-7; mbl-1* which is confirmed in the RT-PCR gel. (See supplemental table 2 for primer sequences).

**Supplemental figure 4.**
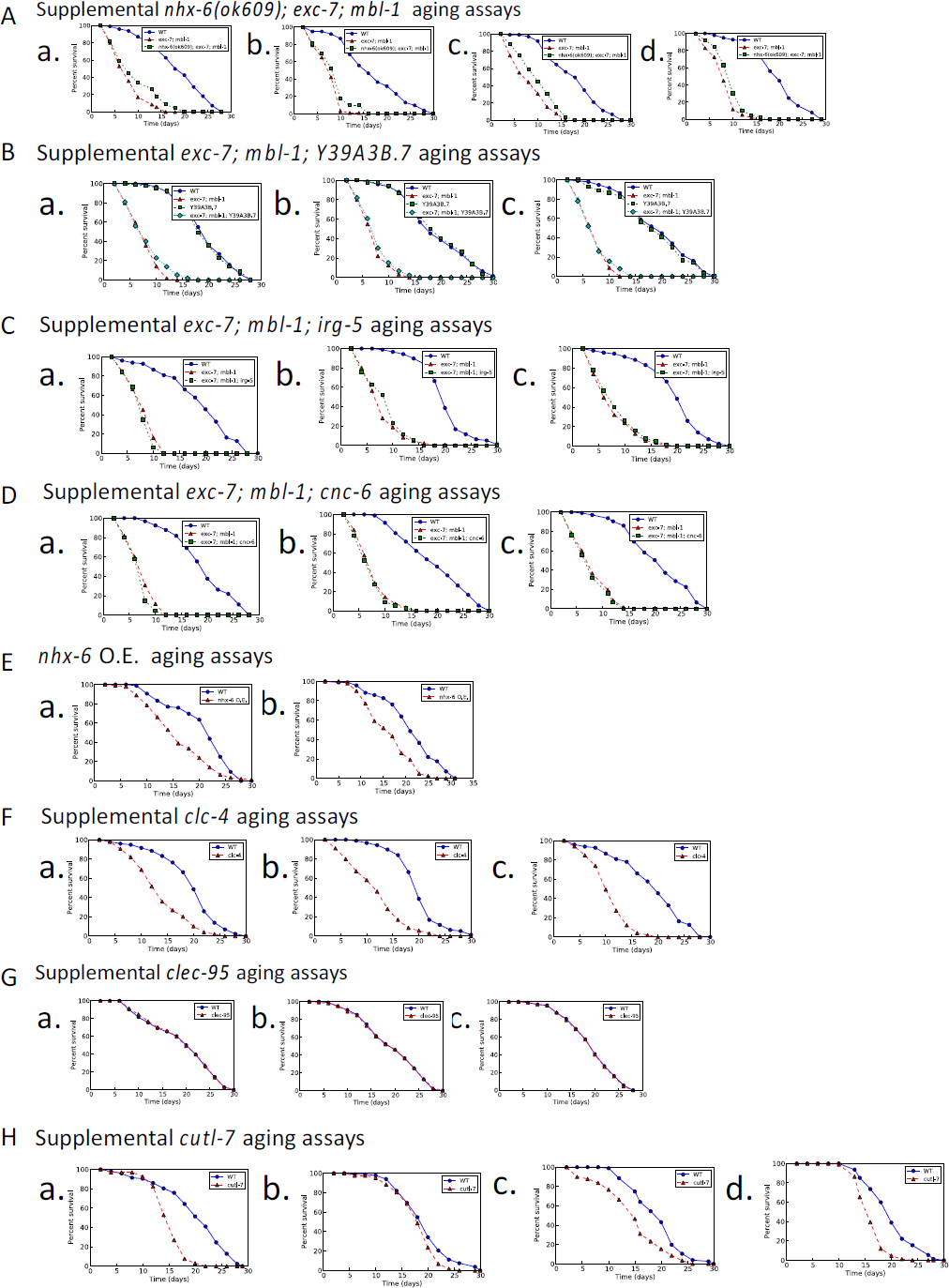
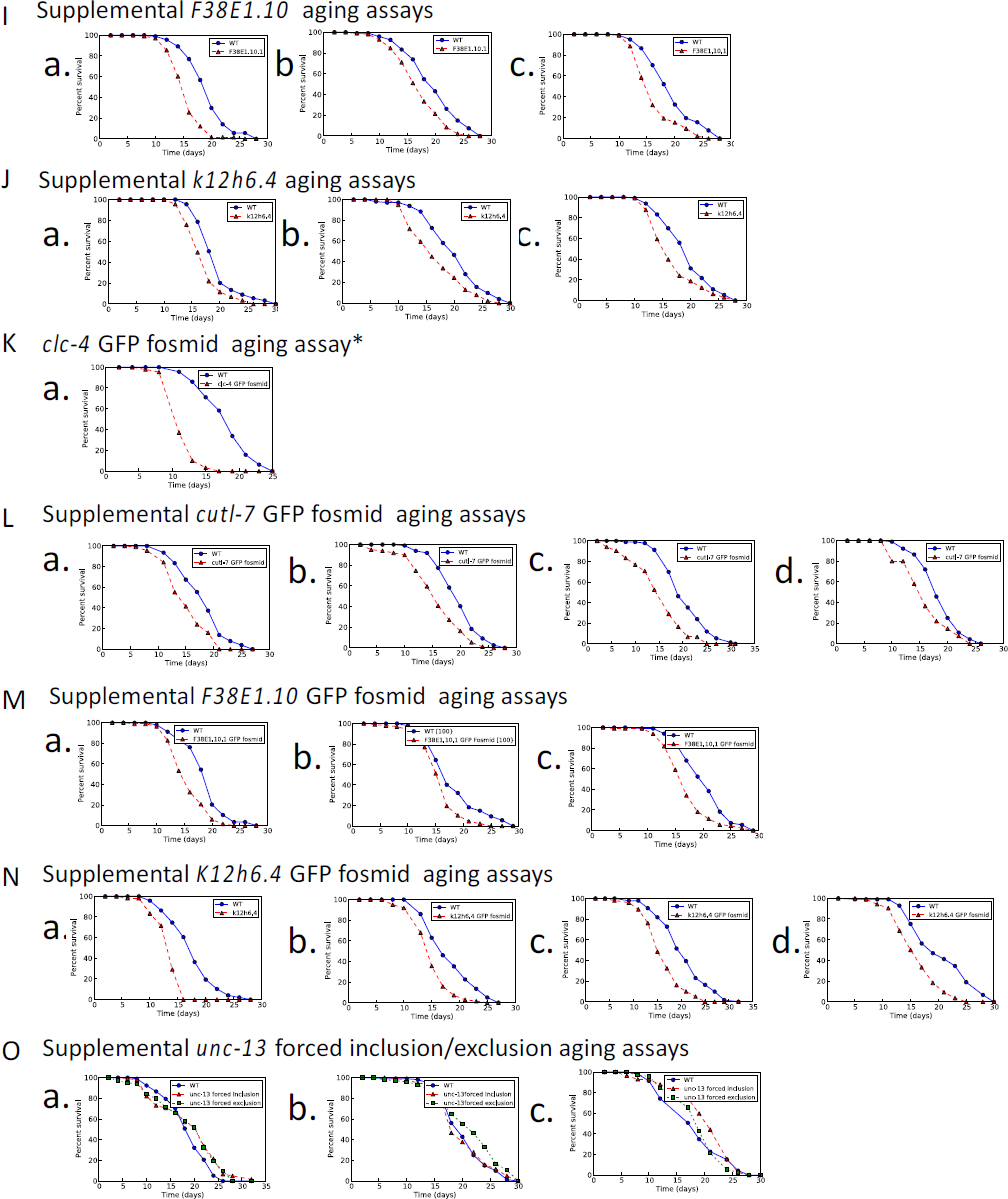
*nhx-6* is partially responsible for *exc-7; mbl-1* lifespan phenotype A. Supplemental *nhx-6(ok609); exc-7; mbl-1* lifespan assays. Bonferroni P-values: a. wild type vs *exc-7; mbl-1=* 0.0e+00, wild type vs *nhx-6(ok609); exc-7; mbl-1*= 0.0e+00, *nhx-6(ok609); exc-7; mbl-1* vs *exc-7; mbl-1*= 0.0497. b. wild type vs *exc-7; mbl-1=* 0.0e+00, wild type vs *nhx-6(ok609); exc-7; mbl-1*= 0.0e+00, *nhx-6(ok609); exc-7; mbl-1* vs *exc-7; mbl-1*= 0.0226. c. wild type vs *exc-7; mbl-1=* 0.0e+00, wild type vs *nhx-6(ok609); exc-7; mbl-1*= 0.0e+00, *nhx-6(ok609); exc-7; mbl-1* vs *exc-7; mbl-1*= 0.0100. d. wild type vs *exc-7; mbl-1=* 0.0e+00, wild type vs *nhx-6(ok609); exc-7; mbl-1*= 0.0e+00, *nhx-6(ok609); exc-7; mbl-1* vs *exc-7; mbl-1*= 0.0022. **B. Supplemental *exc-7; mbl-1; Y39A3B.7* lifespan assays.** Bonferroni P-values: a. wild type vs *Y39A3B.7*= 1, wild type vs *exc-7; mbl-1*= 0.00e+00, wild type vs *exc-7; mbl-1; Y39A3B.7* = 0.00e+00, *exc-7; mbl-1* vs *exc-7; mbl-1; Y39A3B.7*= 0.04959. b. wild type vs *Y39A3B.7*= 1, wild type vs *exc-7; mbl-1*= 0.00e+00, wild type vs *exc-7; mbl-1; Y39A3B.7* = 0.00e+00, *exc-7; mbl-1* vs *exc-7; mbl-1; Y39A3B.7*= 0.6557. c. wild type vs *Y39A3B.7*= 1, wild type vs *exc-7; mbl-1*= 0.00e+00, wild type vs *exc-7; mbl-1; Y39A3B.7* = 0.00e+00, *exc-7; mbl-1* vs *exc-7; mbl-1; Y39A3B.7*= 1. **C. Supplemental *exc-7; mbl-1; irg-5* lifespan assays.** Bonferroni P-values: a. wild type vs *exc-7; mbl-1*= 0.0e+00, wild type vs *exc-7; mbl-1; irg-5*= 0.0e+00, *exc-7; mbl-1* vs *exc-7; mbl-1; irg-5*= 0.4827. b. wild type vs *exc-7; mbl-1*= 0.0e+00, wild type vs *exc-7; mbl-1; irg-5*= 0.0e+00, *exc-7; mbl-1* vs *exc-7; mbl-1; irg-5*= 0.3889. c. wild type vs *exc-7; mbl-1*= 0.0e+00, wild type vs *exc-7; mbl-1; irg-5*= 0.0e+00, *exc-7; mbl-1* vs *exc-7; mbl-1; irg-5*= 0.6572. **D. Supplemental *exc-7; mbl-1; cnc-6* lifespan assays.** Bonferroni P-values: a. wild type vs *exc-7; mbl-1*= 0.0e+00, wild type vs *exc-7; mbl-1; cnc-6*= 0.0e+00, *exc-7; mbl-1* vs *exc-7; mbl-1; cnc-6*= 0.4140. b. wild type vs *exc-7; mbl-1*= 0.0e+00, wild type vs *exc-7; mbl-1; cnc-6*= 0.0e+00, *exc-7; mbl-1* vs *exc-7; mbl-1; cnc-6*= 0.9881. c. wild type vs *exc-7; mbl-1*= 0.0e+00, wild type vs *exc-7; mbl-1; cnc-6*= 0.0e+00, *exc-7; mbl-1* vs *exc-7; mbl-1; cnc-6*= 0.8364**. E. Supplemental *nhx-6* O.E. lifespan assays.** Bonferroni P-values: a. wild type vs *nhx-6* O.E.= 7.6e-07. b. wild type vs *nhx-6* O.E.= 8.5e-09**. F. Supplemental *clc-4* lifespan assays.** Bonferroni P-values: a. wild type vs *clc-4*= 0.0e+00. b. wild type vs *clc-4*= 0.0e+00. c. wild type vs *clc-4*= 0.0e+00. **G. Supplemental *clec-95* lifespan assays.** Bonferroni P-values: a. wild type vs *clec-95*= 0.9647. b. wild type vs *clec-95*= 0.9482. c. wild type vs *clec-95*= 0.8408. **H. Supplemental *cutl-7* lifespan assays.** Bonferroni P-values: a. wild type vs *cutl-7*= 0.0e+00. b. wild type vs *cutl-7*= 0.0485. c. wild type vs *cutl-7*= 1.6e-06 d. wild type vs *cutl-7*= 0.0e+00. **I. Supplemental *F38E1.10.1* lifespan assays.** Bonferroni P-values: a. wild type vs *F38E1.10.1* = 0.0e+00. b. wild type vs *F38E1.10.1* = 0.0001. c. wild type vs *F38E1.10.1* = 3.1e-06. **J. Supplemental *k12h6.4* lifespan assay.** Bonferroni P-values: a. wild type vs *k12h6.4* = 2.1e-05. b. wild type vs *k12h6.4* = 0.0001. c. wild type vs *k12h6.4* = 0.0006**. K. *clc-4* GFP fosmid lifespan assay.** Bonferroni P-values: a. wild type vs *clc-4* GFP fosmid = 0.0e+00. **L. Supplemental *cutl-7* GFP fosmid lifespan assays.** Bonferroni P-values: a. wild type vs *cutl-7* GFP fosmid = 1.3e-05. b. wild type vs *cutl-7* GFP fosmid = 6.5e-08. c. wild type vs *cutl-7* GFP fosmid = 0.0e+00 d. wild type vs *cutl-7* GFP fosmid = 0.0218. **M. Supplemental *F38E1.10.1* lifespan assays.** Bonferroni P-values: a. wild type vs *F38E1.10.1* = 4.1e-07. b. wild type vs *F38E1.10.1* = 0.0002. c. wild type vs *F38E1.10.1* = 1.5e-05. **N. Supplemental *k12h6.4* lifespan assays.** Bonferroni P-values: a. wild type vs *k12h6.4* = 1.6e-08. b. wild type vs *k12h6.4* = 2.1e-07. c. wild type vs *k12h6.4* = 1.0e-08. d. wild type vs *k12h6.4* = 4.1e-08. **O. Supplemental *unc-13* forced inclusion/exclusion lifespan assays.** Bonferroni P-values: a. wild type vs *unc-13* forced inclusion = 0.2165, wild type vs *unc-13* forced exclusion = 0.2457, *unc-13* forced inclusion vs *unc-13* forced exclusion= 1. b. wild type vs *unc-13* forced inclusion = 1, wild type vs *unc-13* forced exclusion = 0.0295, *unc-13* forced inclusion vs *unc-13* forced exclusion= 0.0766. c. wild type vs *unc-13* forced inclusion = 0.0357, wild type vs *unc-13* forced exclusion = 0.8596, *unc-13* forced inclusion vs *unc-13* forced exclusion= 0.0406.

**Supplemental Figure 5.**
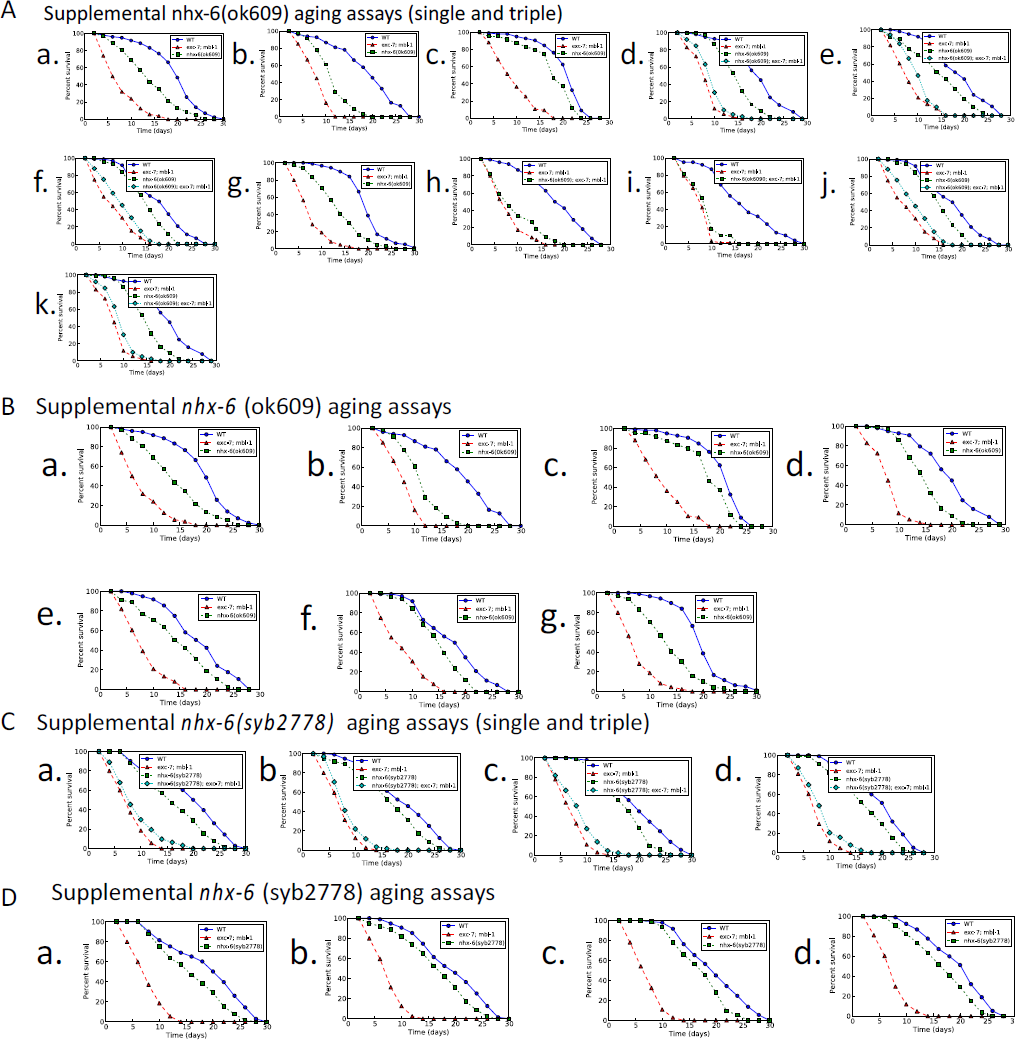
A. Supplemental *nhx-6(ok609)* lifespan assays (single and triple). Bonferroni P-values: a. wild type vs *exc-7;* mbl-1= 0.0e+00, wild type vs *nhx-6(ok609*)= 1.3e-08, *exc-7; mbl-1* vs *nhx-6(ok609*)= 0.0e+00. b. wild type vs *exc-7;* mbl-1= 0.0e+00, wild type vs *nhx-6(ok609*)= 0.0e+00, *exc-7; mbl-1* vs *nhx-6(ok609*)= 6.0e-09 . c. wild type vs *exc-7;* mbl-1= 0.0e+00, wild type vs *nhx-6(ok609*)= 0.0002, *exc-7; mbl-1* vs *nhx-6(ok609*)= 0.0e+00. d. wild type vs *exc-7;* mbl-1= 0.0e+00, wild type vs *nhx-6(ok609*)= 0.0e+00, wild type vs *nhx-6(ok609); exc-7; mbl-1*=0.0e+00, exc-7*; mbl-1* vs *nhx-6(ok609*)= 0.0e+00, *exc-7; mbl-1* vs *nhx-6(ok609); exc-7; mbl-1*= 0.0033. e. wild type vs *exc-7;* mbl-1= 0.0e+00, wild type vs *nhx-6(ok609*)= 0.0002, wild type vs *nhx-6(ok609); exc-7; mbl-1*=0.0e+00, *exc-7; mbl-1* vs *nhx-6(ok609*)= 0.0e+00, *exc-7; mbl-1* vs *nhx-6(ok609); exc-7; mbl-1*= 0.0011. f. wild type vs *exc-7;* mbl-1= 0.0e+00, wild type vs *nhx-6(ok609*)= 0.0009, wild type vs *nhx-6(ok609); exc-7; mbl-1*= 0.0e+00, exc-7*; mbl-1* vs *nhx-6(ok609)*= 0.0e+00, *exc-7; mbl-1* vs *nhx-6(ok609); exc-7; mbl-1*= 0.0150. g. wild type vs *exc-7; mbl-1*= 0.0e+00, wild type vs *nhx-6(ok609*)= 0.0e+00, *exc-7; mbl-1* vs *nhx-6(ok609*)= 0.0e+00. h. wild type *vs exc-7; mbl-1*= 0.0e+00, wild type vs *nhx-6(ok609*); *exc-7; mbl-1*= 0.0e+00, *exc-7; mbl-1* vs *nhx-6(ok609); exc-7; mbl-1*= 0.0497. i. wild type vs *exc-7;* mbl-1= 0.0e+00, wild type vs *nhx-6(ok609*); *exc-7; mbl-1*= 0.0e+00, *exc-7; mbl-1* vs *nhx-6(ok609*); *exc-7; mbl-1*= 0.0226. j. wild type vs *exc-7;* mbl-1= 0.0e+00, wild type vs *nhx-6(ok609*)= 0.0009, wild type vs *nhx-6(ok609); exc-7; mbl-1*=0.0e+00, exc-7*; mbl-1* vs *nhx-6(ok609*)= 0.0e+00, *exc-7; mbl-1* vs *nhx-6(ok609); exc-7; mbl-1*= 0.0150. k. wild type vs *exc-7;* mbl-1= 0.0e+00, wild type vs *nhx-6(ok609*)= 0.0e+00, wild type vs *nhx-6(ok609); exc-7; mbl-1*=0.0e+00, exc-7*; mbl-1* vs *nhx-6(ok609*)= 0.0e+00, *exc-7; mbl-1* vs *nhx-6(ok609); exc-7; mbl-1*= 0.0033. **B. Supplemental *nhx-6(ok609)* lifespan assays.** Bonferroni P-values: a. wild type vs *exc-7; mbl-1*= 0.0e+00, wild type vs *nhx-6(ok609)*= 1.3e-08, *exc-7; mbl-1* vs *nhx-6(ok609)*= 0.0e+00. b. wild type vs *exc-7; mbl-1*= 0.0e+00, wild type vs *nhx-6(ok609)*= 0.0e+00, *exc-7; mbl-1* vs *nhx-6(ok609)*= 6.0e-09. c. wild type vs *exc-7; mbl-1*= 0.0e+00, wild type vs *nhx-6(ok609)*= 0.00e+00, *exc-7; mbl-1* vs *nhx-6(ok609)*= 0.0002 d. wild type vs *exc-7; mbl-1*= 0.0e+00, wild type vs *nhx-6(ok609)*= 0.0e+00. e. wild type vs *exc-7; mbl-1*= 0.0e+00, wild type vs *nhx-6(ok609)*= 0.0001, *exc-7; mbl-1* vs *nhx-6(ok609)*= 0.0e+00. f. wild type vs *exc-7; mbl-1*= 0.0e+00, wild type vs *nhx-6(ok609)*= 0.0006, *exc-7; mbl-1* vs *nhx-6(ok609)*= 0.0e+00. g. wild type vs *exc-7; mbl-1*= 0.0e+00, wild type vs *nhx-6(ok609)*= 0.0e+00, *exc-7; mbl-1* vs *nhx-6(ok609)*= 0.0e+00. **C. Supplemental *nhx-6(syb2778)* lifespan assays (single and triple).** Bonferroni P-values: a. wild type vs *exc-7; mbl-1*= 0.0e+00, wild type vs *nhx-6(syb2778)*= 0.0012, wild type vs *nhx-6(syb2778); exc-7; mbl-1*= 0.0e+00, *exc-7, mbl-1* vs *nhx-6(syb2778); exc-7; mbl-1*= 0.0564. b. wild type vs *exc-7; mbl-1*= 0.0e+00, wild type vs *nhx-6(syb2778)*= 0.0106, wild type vs *nhx-6(syb2778); exc-7; mbl-1*= 0.0e+00, *exc-7, mbl-1* vs *nhx-6(syb2778); exc-7; mbl-1*= 0.0230. c. wild type vs *exc-7; mbl-1*= 0.0e+00, wild type vs *nhx-6(syb2778)*= 0.0014, wild type vs *nhx-6(syb2778); exc-7; mbl-1*= 0.0e+00, *exc-7, mbl-1* vs *nhx-6(syb2778); exc-7; mbl-1*= 0.0050. d. wild type vs *exc-7; mbl-1*= 0.0e+00, wild type vs *nhx-6(syb2778)*= 0.0011, wild type vs *nhx-6(syb2778); exc-7; mbl-1*= 0.0e+00, *exc-7, mbl-1* vs *nhx-6(syb2778); exc-7; mbl-1*= 0.0251. **D. Supplemental *nhx-6(syb2778)* lifespan assays.** Bonferroni P-values: a. wild type vs *exc-7; mbl-1*= 0.0e+00, wild type vs *nhx-6(syb2778)*= 0.0008, e*xc-7; mbl-1* vs *nhx-6(syb2778)*= 0.0e+00. b. wild type vs *exc-7; mbl-1*= 0.0e+00, wild type vs *nhx-6(syb2778)*= 0.0071, e*xc-7; mbl-1* vs *nhx-6(syb2778)*= 0.0e+00. c. wild type vs *exc-7; mbl-1*= 0.0e+00, wild type vs *nhx-6(syb2778)*= 0.0009, e*xc-7; mbl-1* vs *nhx-6(syb2778)*= 0.0e+00, d. wild type vs *exc-7; mbl-1*= 0.0e+00, wild type vs *nhx-6(syb2778)*= 0.0007, e*xc-7; mbl-1* vs *nhx-6(syb2778)*= 0.0e+00

